# Isolation of postnatal human neural stem cells

**DOI:** 10.64898/2026.06.11.731596

**Authors:** Daniel Dan Liu, Anna E. Eastman, Nicole L. Womack-Gambrel, Chang N. Kim, Joy Q. He, Suyash Raj, Emily Reilly, Rahul Sinha, Nobuko Uchida, Kelly Chau, Benjamin F. Ohene-Gambill, Samrat Thapa, Emon Nasajpour, Julia A. Belk, Norma F. Neff, Siddhartha Jaiswal, H. Westley Phillips, Meagan Chambers, Claudia K. Petritsch, Gerald A. Grant, Laura M. Prolo, Jody E. Hooper, Tomasz J. Nowakowski, Irving L. Weissman

## Abstract

While it was once thought that neurogenesis is complete by birth, it is now apparent that the human brain continues to generate new neurons postnatally, at least into childhood. While much attention has been focused on postnatally-born neurons, their presumed progenitor – the postnatal neural stem cell (NSC) – remains poorly characterized. Using index sorting, we identify and prospectively isolate two subsets of NSCs from the postnatal human brain, and describe their differentiation dynamics using clonal barcoding and in vivo xenotransplantation. We demonstrate an A2B5^+^EGFR^+^ population biased towards interneuron and oligodendrocyte fates (NINO), and an A2B5^−^EGFR^hi^ population biased towards an astrocyte fate (NAC). Profiling of human brains across lifespan shows that the frequency of NSCs declined exponentially across the first two decades of life, but stabilized thereafter, still present in the brains of donors as old as 90 years. Our study provides a framework for the functional study of postnatal human NSCs and their potential roles in development, aging, and disease.

## INTRODUCTION

Neural stem cells (NSCs) represent a highly specialized tissue resident stem cell population that self-renews and differentiates to produce the neurons, oligodendrocytes, astrocytes, and ependymal cells of the brain. Across species, the majority of these cells are found during early embryonic development, and take the form of radial glia that reside near the ventricular zone (VZ), and produce cells in a temporally- and spatially-ordered manner^1,2^. Near the end of gestation and entering postnatal life, the vast majority of NSCs enter quiescence or differentiate into astrocytes^3–5^. It was once assumed that the adult brain produced no new neurons^6^, but this dogma has been disproven through multiple experimental paradigms: initially in the 1960s with radioisotope labeling experiments in canaries^7^, rats^8,9^, and cats^10^, then later in the 1990s by the isolation and in vitro culture of neural cells with multilineage differentiation capabilities from mice^11–13^ and rats^14,15^, BrdU labeling in mice^16^, rats^17^, macaques^18^, and humans^19^, and lineage tracing and grafting experiments in mice^20–22^. Two adult NSC niches are widely recognized^23^: first, the ventricular-subventricular zone (V-SVZ) on the walls of the lateral ventricles (LV), where NSCs called B1 cells generate interneurons that migrate through the rostral migratory stream into the olfactory bulb^20,21^; second, the subgranular zone (SGZ) of the hippocampus, where NSCs variously called radial glia-like cells (RGLs), radial astrocytes, or type 1 progenitors, generate new excitatory neurons for the dentate gyrus (DG)^15–18,22^.

It remains controversial whether the results of these experiments, mostly performed in rodents, hold true in the human brain as well. Regarding the hippocampus, many studies have reported supportive evidence^19,24–29^, or lack thereof^30–33^, for human hippocampal neurogenesis beyond childhood. Setting adulthood aside, it is less controversial that in infancy, several streams of newborn interneurons persist that deliver neurons to specific brain regions^34^. These include the rostral and medial migratory streams (RMS, MMS) delivering neurons to the olfactory bulb and ventromedial prefrontal cortex, respectively^35^, the Arc delivering neurons to the cingulate and superior frontal gyri^36^, and the entorhinal cortex stream^37^. Recent studies also suggest that excitatory neurons may still be born in late development or early infancy^37,38^, though their source and ultimate fate are still to be worked out.

While many studies have focused on the nascent neurons, less is understood about their origin, the postnatal NSC. Previous studies have generally utilized bulk culture of micro-dissected human brain without purification^39,40^, or sorting on cell-type-specific promoter-driven GFP expression^41,42^. We previously described the prospective isolation of ten neural stem and progenitor cell (NSPC) types from second trimester human brain tissue^43–45^, from the radial glia to the mature neurons and glia. Here, we again combine flow cytometry with single cell transcriptomics to purify and functionally interrogate NSCs, now from the postnatal human brain. We utilize index-sorting to map single cell transcriptomes to their surface marker expression (immunophenotype), deriving a purification strategy. Our gating scheme allows for the prospective isolation of two subsets of postnatal NSC – one biased towards interneurons and oligodendrocyte precursor cells (OPCs), and the other towards astrocytes – in addition to simultaneous purification of OPCs, oligodendrocytes, astrocytes, and non-neural lineages. Our findings establish a framework for the functional interrogation of postnatal human NSCs, and for future studies on their role in postnatal neurodevelopment, stem cell aging, and neuropsychiatric disorders.

## RESULTS

### Identification of a neural stem cell-like population from postnatal human brain

We utilized two types of viable human tissue for this study: neurosurgical resections and rapid autopsy **(Supplementary Table 1)**. Neurosurgical tissue has the advantage of higher viability, with the tradeoff that the tissue necessarily has some type of pathology, typically epilepsy (though any tissues with known neoplasms were excluded for functional experiments). Rapid autopsy tissue has the advantage of patient selection to exclude those with known neoplastic disease, but typically comprises older donors, and has a longer postmortem interval (3-15 hr).

To identify candidate neural stem cell (NSC) populations, we dissociated postnatal human brain tissue into a single-cell suspension **(Fig. 1a)**. Notably, postnatal brain tissue necessitated a papain-based dissociation for optimal cell recovery and viability, which unlike our protocol for prenatal brain tissue^43,44^, results in the proteolytic cleavage of many cell surface markers. Cells were stained with a panel of antibodies against EGFR, A2B5, CD90 (THY1), and non-neural lineages (CD45 [PTPRC] for hematopoietic lineages; CD31 [PECAM1], CD34, and CD105 [ENG] for vascular lineages; CD235a [GYPA] for erythrocytes), then analyzed using flow cytometry. After pre-gating for live neural-lineage singlets, we identified A2B5 and EGFR as additional markers that split the neural compartment **(Fig. 1b, Extended Data Fig. 1a)**. While most neural cells were A2B5^−^EGFR^−^, we also identified single-positive A2B5^+^EGFR^−^, A2B5^−^EGFR^mid^, and A2B5^−^EGFR^hi^ cells, as well as rare double-positive A2B5^+^EGFR^+^ cells. To determine the identity of these populations, we used fluorescence-activated cell sorting (FACS) to index-sort cells for single-cell RNA sequencing (scRNA-seq) with Smart-seq3^46^. Index sort records the mean fluorescence intensity (MFI) of each surface marker for each sorted single cell, which, due to the plate-based format of Smart-seq3, allows for the integration of transcriptomic and immunophenotypic identity^43^. A total of 4 donors were sequenced: a 9-day-old male born at term who passed from apnea, two 9-month-old females who underwent hemispherectomy for hemimegaloencephaly (HME), and one 9-year old male who underwent surgical resection for epilepsy secondary to tuberous sclerosis (TSC).

**Figure 1:**
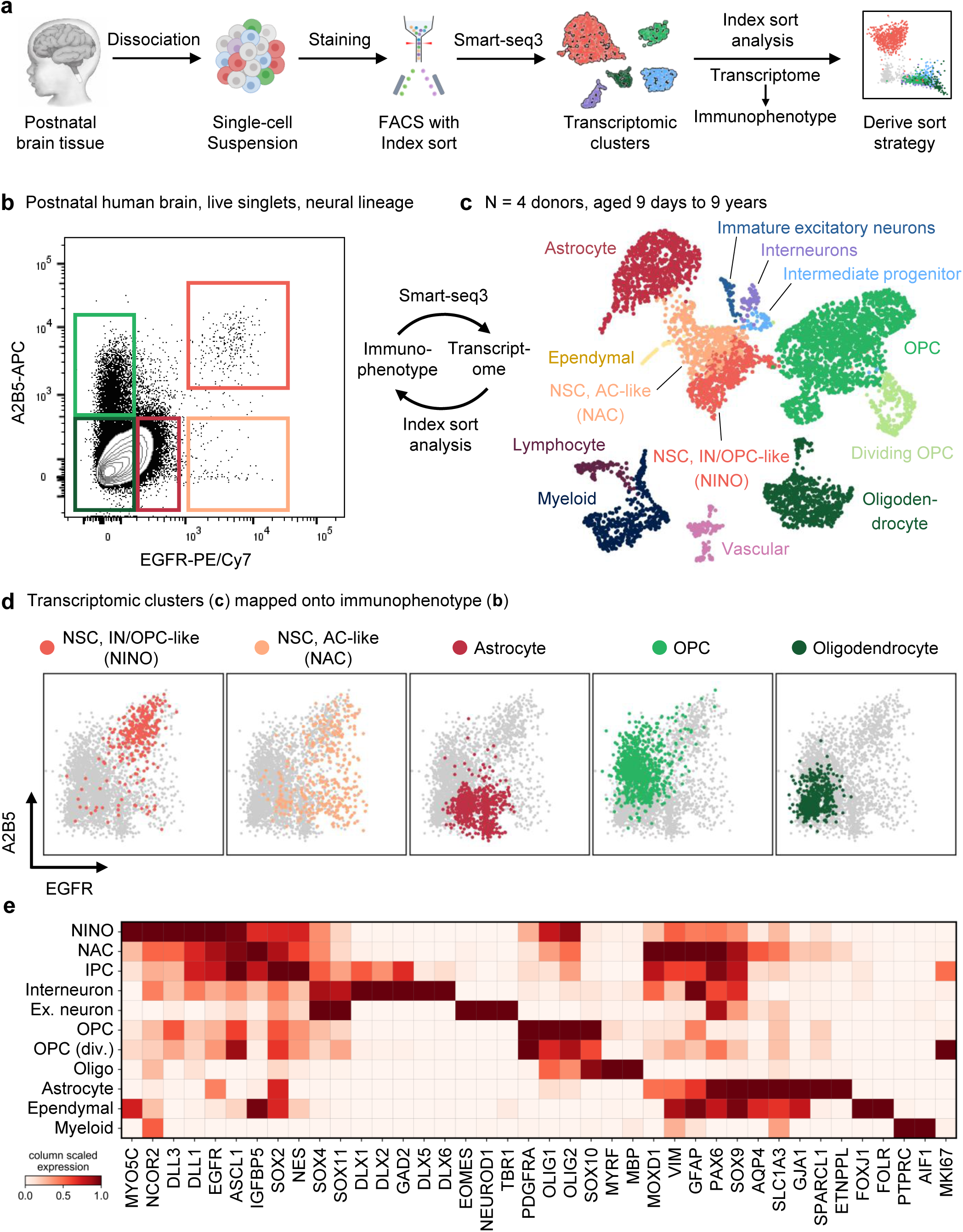
Prospective isolation of a putative postnatal human neural stem cell. **a,** Experimental workflow used to derive purification strategy for postnatal human neural stem cells. **b,** Flow cytometry gating for purifying postnatal human neural cells. Events are pregated on live singlets negative for non-neural linages (CD45, CD11b, CD105, CD31, CD34). **c,** UMAP plot with cluster annotations of postnatal human brain cells sequenced using Smart-seq3. **d,** Index sort analysis mapping transcriptomically-defined clusters onto their surface marker profile (immunophenotype). **e,** Matrix plot showing expression of marker genes by transcriptomic cluster from primary postnatal human brain cells.

Transcriptomic analysis identified known neural cell types based on standard marker genes. These include astrocytes (*GFAP*, *AQP4*), oligodendrocyte precursor cells (OPCs) (*OLIG2*, *SOX10*, *PDGFRA*), oligodendrocytes (*SOX10*, *MYRF*, and *MBP*), and ependymal cells (*FOXJ1*, *FOLR1*) **(Fig. 1c, d)**. Non-neural cell types such as myeloid, lymphoid, and vascular cells were also captured for internal comparison. In addition to these expected cell types, we identified a distinct cluster of cells expressing NSC-associated genes such as *NES*, *EGFR*, and *ASCL1*. Also present were a small number of dividing intermediate progenitor cells (IPCs) likely giving rise to inhibitory neurons, based on their expression of DLX transcription factors and *GAD2*. Finally, we captured a small number of immature excitatory neurons expressing *NEUROD1* and deep-layer marker *TBR1*. Immature neurons were present in 3 of the 4 donors, with none recovered from the 9-year-old donor **(Extended Data Fig. 1b, c)**. These findings are consistent with recent reports of protracted generation of excitatory and inhibitory neurons into early postnatal life^37,38^.

Further sub-clustering of the putative NSC cluster identified two main subsets. The first subset expressed astrocyte-associated genes (such as *SOX9*, *AQP4*, *APOE*, and *FAM107A*), though lacked other mature astrocyte markers (such as *SPARCL1* and *ETNPPL*) **(Fig. 1c, d; Extended Data Fig. 2)**. The second subset expressed a mixture of OPC-associated genes (such as *PDGFRA* and *OLIG1/2*), neuron-associated genes (such as *DLL1/3* and *DCX*), as well as more specific genes (such as *MYO5C* and *NCOR2*), though lacked more lineage-committed markers (such as *SOX10* or *DLX* transcription factors). Both subsets expressed common genes such as *EGFR*, *ASCL1*, *NES*, and *BTG2*. Both NSC subsets were present in all 4 donors **(Extended Data Fig. 1b, c)**. We will subsequently refer to the former subpopulation as “NACs” (<u>N</u>SC-<u>a</u>stro<u>c</u>yte-like) and the latter as “NINOs” (<u>N</u>SC-inter<u>n</u>euron/<u>O</u>PC-like).

Histological analysis confirmed the presence of cells co-expressing NES and *EGFR* transcript in human subventricular zone (SVZ) **(Extended Data Fig. 3a)**. Further staining demonstrated heterogeneity among NES-expressing cells in the SVZ, with some expressing high levels of PDGFRA and low SOX9, consistent with a NINO identity, and others expressing high levels of SOX9 and low PDGFRA, consistent with a NAC identity **(Extended Data Fig. 3b)**. We did observe some NES-expressing cells that co-expressed both PDGFRA and SOX9, suggesting that the NINO-NAC distinction may be more a continuum than a binary.

As powerful as transcriptomic analysis is for cell type identification, stem cells are ultimately defined by their functional properties of self-renewal and multipotency—purification of our putative NSCs is therefore requisite for experimental tractability. To this end, we utilized index sort analysis, which maps transcriptomically defined clusters onto their surface marker profile (immunophenotype). Index sort analysis revealed that the neural cell types within our dataset segregated based on surface expression of A2B5 and EGFR **(Fig. 1e; Extended Data Fig. 4a)**. OPCs were A2B5^+^EGFR^−^; oligodendrocytes were A2B5^−^EGFR^−^, and astrocytes were A2B5^−^EGFR^mid^. Importantly, the NINO population was uniquely A2B5^+^EGFR^+^, while the NAC population was largely A2B5^−^EGFR^hi^. As an internal control, we found that as expected, myeloid cells were CD45^+^CD11b^+^ and lymphocytes CD45^+^CD11b^−^ **(Extended Data Fig. 4b)**. These results were consistent across all four donors **(Extended Data Fig. 5)**. Unique immunophenotypes allow for the quantification and, more importantly, the purification of our candidate NSC populations.

### Postnatal human NSCs are tripotent in vitro

Though the transcriptomic and histologic characteristics are supportive of a stem cell identity, the definition of a stem cell is inherently a functional one, defined by self-renewal and multilineage differentiation. We thus purified A2B5^+^EGFR^+^ (enriched for NINOs) and A2B5^−^EGFR^hi^ (enriched for NACs) from postnatal human brain tissue for in vitro culture **(Fig. 2a)**. Immunocytochemistry showed that purified A2B5^+^EGFR^+^ cells uniformly express Nestin, with a subset also co-expressing GFAP **(Extended Data Fig. 6a)**. These cells were capable of serial passage in media containing EGF and FGF, upon which they again give rise to Nestin^+^ colonies of cells, demonstrating self-renewal capability. In contrast, purified A2B5^+^EGFR^−^ cells expressed O4, consistent with an oligodendrocyte identity, while purified A2B5^−^EGFR^mid^ cells expressed GFAP but not SOX2, consistent with an astrocyte identity **(Extended Data Fig. 6b)**, neither of which demonstrated proliferative potential.

**Figure 2:**
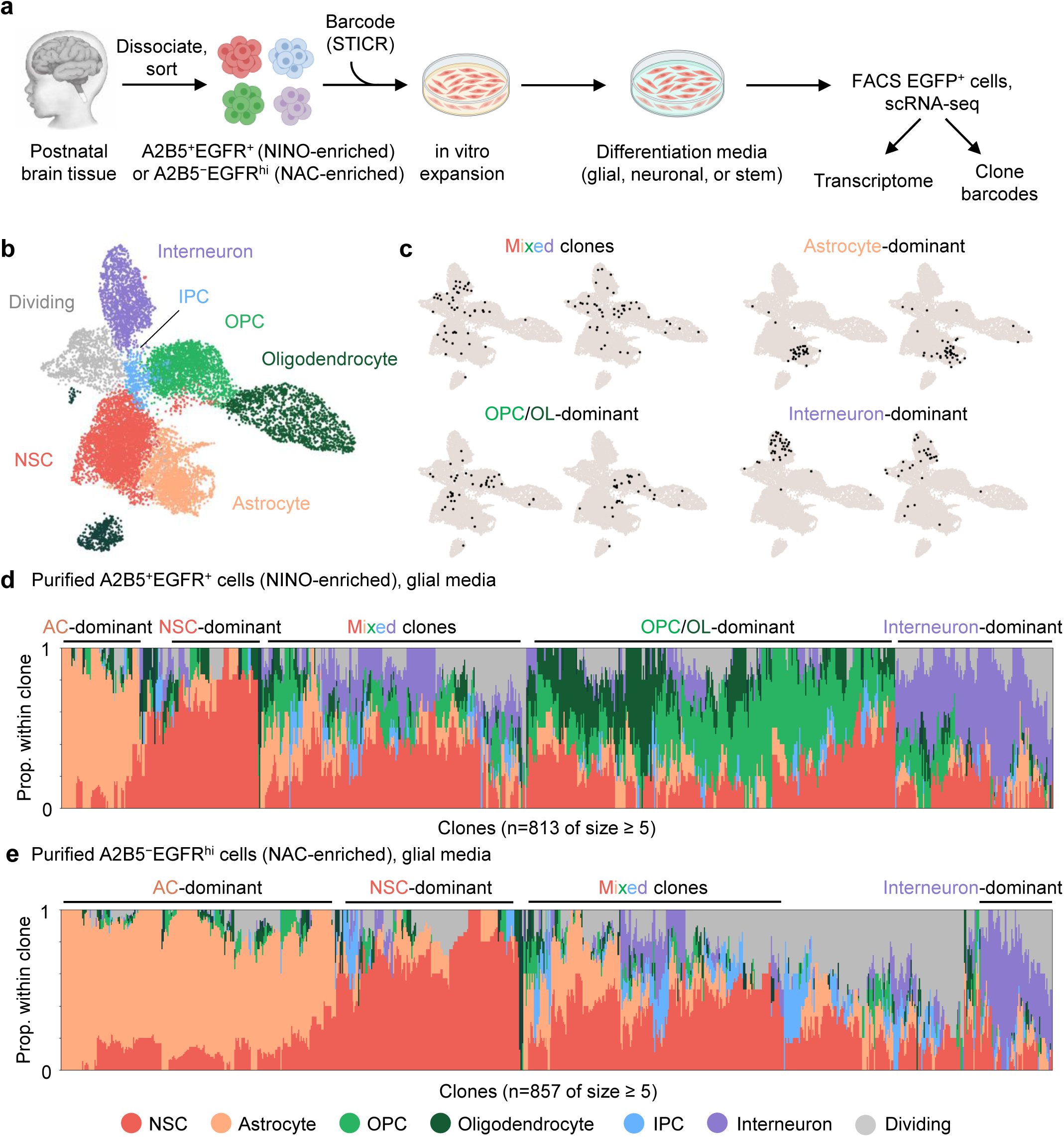
Putative NSCs are tripotent in vitro. **a,** Experimental workflow used for in vitro barcoding experiments. **b,** UMAP plot with cluster annotations of purified postnatal human NSCs following in vitro culture. **c,** Selected clones recovered from barcode sequencing, imposed on UMAP. Black dots within a plot indicate cells with shared barcodes. **d–e,** Stacked bar plots showing relative cell type makeup within each clone (size ≥ 5) recovered following culture of A2B5^+^EGFR^+^ cells (**d**) or A2B5^−^EGFR^hi^ cells (**e**). Each column represents one clone.

Next, we sought to functionally interrogate the differentiation potential of the putative NSCs. To track cells at a clonal level, we introduced unique genetic barcodes via a lentivirus shortly after cell sorting^47^. For the following experiments, we utilized cells derived from the SVZ of a 9-day-old male donor, born at term, who passed from apnea. Cells were allowed to expand for 7 days, and were then put in one of 3 culture medias: (1) stem cell media containing EGF and FGF, (2) glial media containing PDGF, NT3, and IGF, or (3) neuron media containing BDNF and GDNF. Following 5 days of differentiation, cells were sorted for scRNA-seq for joint transcriptomic and clonal barcode readout.

Single cell transcriptomic analysis demonstrated that cultured NSCs self-renewed and differentiated into all 3 major neural lineages: oligodendrocytes, astrocytes, and interneurons **(Fig. 2b, Extended Data Fig. 6c)**. Interneurons expressed *NR2F1/2* and *MEIS2*, but no *NKX2.1* or *LHX6*, suggesting a caudal ganglionic eminence (CGE) rather than medial ganglionic eminence (MGE) origin. They furthermore expressed *TAC3*, *CALB2*, and *NPY*, but not *VIP* and little *SST* **(Extended Data Fig. 6d)**.

The relative proportion of these cell types varied based on the sorted population and the culture media **(Extended Data Fig. 7a)**. Comparing A2B5^+^EGFR^+^ cells (enriched for NINOs) versus A2B5^−^EGFR^hi^ cells (enriched for NACs), the former produced a notably higher proportion of oligodendroglia, while the latter produced a higher proportion of astrocytes, in all 3 media conditions. In both A2B5^+^EGFR^+^ and A2B5^−^EGFR^hi^ populations, culturing in neuron media increased the relative proportion of neurons, while decreasing the proportion of NSCs.

Clonal barcode analysis recovered a total of 2,369 clones of size 5 or greater. Clones were identified spanning all neural lineages: NSC, interneuron, oligodendrocyte, and astrocyte **(Fig. 2c)**, demonstrating that our purified cells have both self-renewal and trilineage differentiation potential. Certain clones showed strong lineage bias towards specific fates, defined here as having >40% of cells within the clone being made up of a single lineage **(Fig. 2d, e; Extended Data Fig. 7b, c)**. Within the glial media condition, A2B5^+^EGFR^+^-derived clones had a sizeable proportion of oligodendrocyte-dominant clones (29.4%) and interneuron-dominant clones (10.8%), and a smaller of astrocyte-dominant clones (9.1%). In contrast, A2B5^−^EGFR^hi^-derived clones had nearly no oligodendrocyte-dominant clones (0.7%), a small proportion of interneuron-dominant clones (4.2%), but a large proportion of astrocyte-dominant clones (29.8%). Culture media also had a strong effect on clone types **(Extended Data Fig. 7b, c)**. For the A2B5^+^EGFR^+^-derived clones, stem cell media resulted in the highest proportion of NSC-dominant clones (43.4%) and astrocyte-dominant clones (20.2%), glial media the highest proportion of oligodendrocyte-dominant clones (29.4%), and neuron media the highest proportion of interneuron-dominant clones (31.3%).

### Postnatal human NSCs engraft and differentiate in vivo

We next turned to an in vivo xenotransplantation model to further support the tripotency of our postnatal NSCs **(Fig. 3a)**. Due to their rarity, purified A2B5^+^EGFR^+^ and A2B5^−^EGFR^hi^ cells were expanded in vitro to obtain sufficient numbers for transplant. Cells were labelled with an EGFP-carrying lentivirus prior to transplantation to facilitate re-isolation. A total of 10^5^ cells were then transplanted into the lateral ventricles of neonatal NOD-*scid*-IL2Rg^null^ (NSG) immunodeficient mice at postnatal day 1. Mice were sacrificed 6 months post-transplant, after which one cerebral hemisphere was fixed for histology, and the other was immediately dissociated to re-isolate engrafted human cells for scRNA-seq.

**Figure 3.**
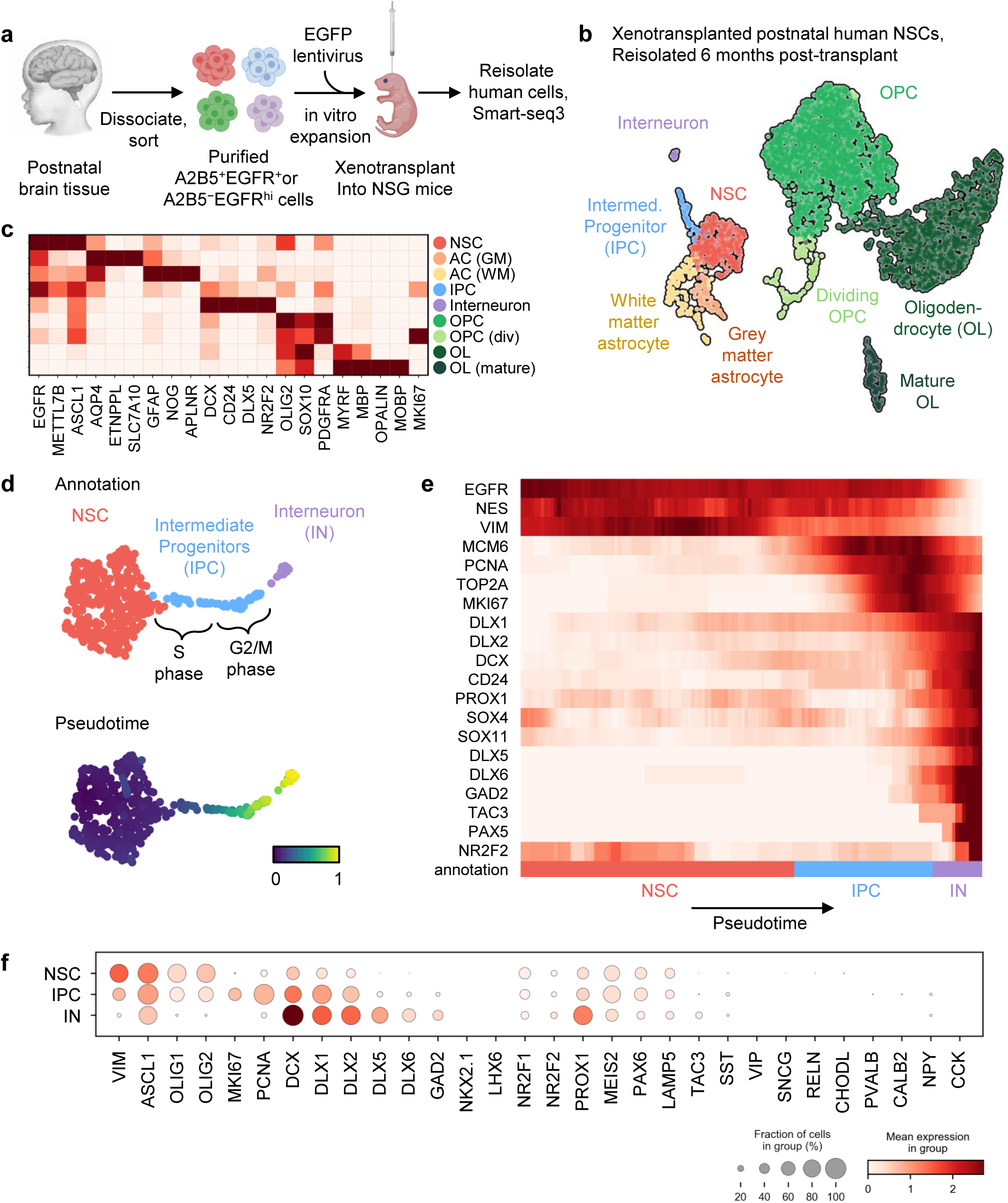
Postnatal human NSCs engraft, self-renew, and differentiate in vivo. **a,** Experimental workflow used for xenotransplant experiments. **b,** UMAP plot with cluster annotations of engrafted human cells, 6 months following transplant of postnatal human NSCs. **c,** Matrix plot showing expression of marker genes by transcriptomic clusters from engrafted human cells. **d,** Pseudotime trajectory analysis of engrafted NSCs, IPCs, and interneurons. **e,** Expression of selected marker genes along the pseudotime trajectory from NSC to IPC to interneuron. **f,** Dot plot showing expression of interneuron subtype marker genes.

Dissociated brains of transplanted mice were analyzed using flow cytometry. A distinct population of EGFP^+^ cells was observed in the transplanted mice but not control littermates, confirming engraftment of human cells **(Extended Data Fig. 8a)**. EGFP^+^ cells were then sorted for scRNA-seq using Smart-seq3. Engraftment was observed in all 18 transplanted mice, from which we obtained single-cell transcriptomes of 3,883 engrafted human cells **(Fig. 3b, Extended Data Fig. 8b)**. Among the engrafted cells we observed NSCs as well as differentiated astrocytes, OPCs, oligodendrocytes, and a small population of IPCs giving rise to interneurons **(Fig. 3b, c)**. Among engrafted astrocytes, we were further able to distinguish grey matter astrocytes (marked by *SLC7A10*, *APOE*, and *ATP1A2*) from white matter astrocytes (marked by *PLP1*, *ZNF704*, and *GFAP*) **(Extended Data Fig. 8c)**, consistent with previous transcriptomic studies in the mouse^48^. Most recovered cells were of oligodendrocyte lineage (84.2%), which may either reflect the gliogenic environment of the mouse brain at time of transplant, or their preferential survival following tissue dissociation.

While the number of engrafted interneurons was small (0.93%), we believe this quantity to be artificially diminished by tissue dissociation, which is known to deplete neurons. Pseudotime analysis demonstrated a trajectory from engrafted NSCs, progressing through S phase then G2/M phase IPCs, to mature interneurons **(Fig. 3d, e)**. Engrafted interneurons expressed *NR2F2* (COUPTFII), *MEIS2*, and *PROX1*, but not *NKX2.1* or *LHX6*, suggestive of a CGE rather than MGE origin **(Fig. 3e, f)**, consistent with the in vitro data **(Extended Data Fig. 6d)**. Comparing A2B5^+^EGFR^+^ cells (enriched for NINOs) versus A2B5^−^EGFR^hi^ cells (enriched for NACs), we find the latter produced a higher proportion of astrocytes (4.4% vs. 6.6%, p=0.018) **(Extended Data Fig. 8d)**.

Histology of transplanted mouse brains was performed in parallel with single-cell transcriptomics. Tissue sections were stained with SC121, a human-specific cytoplasmic marker, as well as lineage-specific markers. The human identity of the engrafted cells could be further confirmed based on their distinct nuclear morphology, with human cells having more diffuse heterochromatin with fewer prominent foci on DAPI staining **(Extended Data Fig. 9a)**. Human cells were found engrafted throughout the mouse brain. SC121^+^OLIG2^+^ OPCs and oligodendrocytes were identified throughout the brain parenchyma and white matter tracts **(Fig. 4a-iii, b)**. SC121^+^GFAP^+^ astrocytes were present **(Fig. 4a-iv, c)** and could occasionally be observed forming end foot processes along blood vessels **(Extended Data Fig. 9b)**, suggesting functional integration. Rare SC121^+^NeuN^+^ neurons were observed in the hippocampus (though not in the dentate gyrus) **(Fig. 4a-i,ii)**. Finally, SC121^+^NES^+^EGFR^+^ cells were observed lining the lateral ventricles, with fine processes extending laterally, likely representing NSCs engrafted in a site-appropriate manner **(Fig. 4d)**.

**Figure 4.**
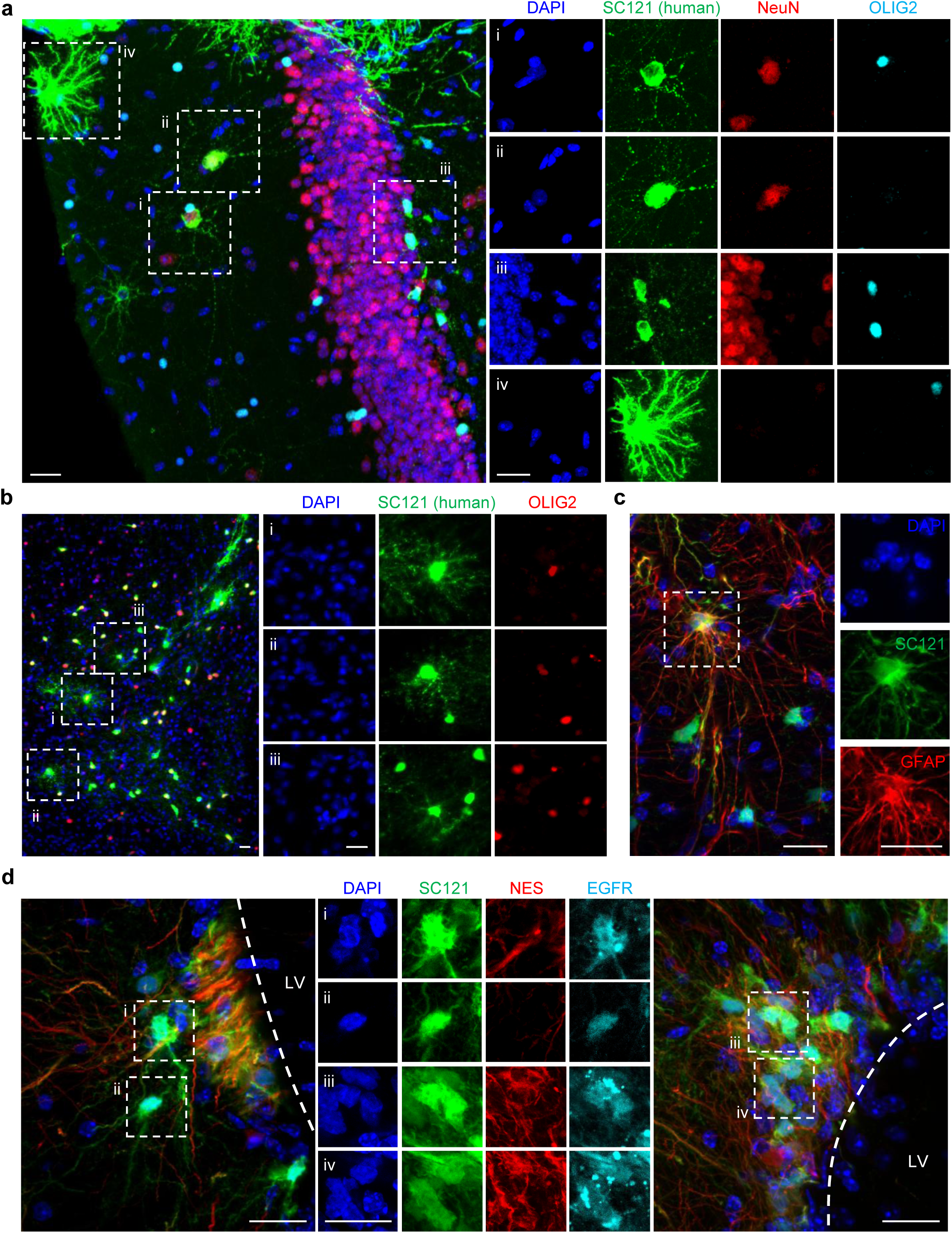
Histology of engrafted postnatal human neural stem cells. **a,** Mouse hippocampus, stained for DAPI (blue), SC121 human cytoplasmic antigen (green), NeuN (red), and OLIG2 (cyan). **b,** Mouse subcortical white matter, stained for DAPI (blue), SC121 (green), and OLIG2 (red). **c,** Mouse subcortical white matter, stained for DAPI (blue), SC121 (green), and OLIG2 (red). **d,** Mouse subventricular zone, stained for DAPI (blue), SC121 (green), NES (red), and EGFR (cyan). LV, lateral ventricle. Scale bars 20 μm.

### A2B5^+^EGFR^+^ cells decrease but persist across lifespan

We next sought to characterize the frequency of postnatal NSCs between brain regions and across lifespan. Viable brain tissue was dissociated and analyzed using flow cytometry from 26 donors, ranging from 9 days to 90 years of age **(Supplementary Table 1)**. We focused on the A2B5^+^EGFR^+^ population as it was more consistent across donors. We first sought to assess whether A2B5^+^EGFR^+^ cells are enriched in classically neurogenic zones, such as the SVZ and hippocampus. This comparison was possible in cases where we received tissue from multiple known brain regions within the same donor. In neurosurgical tissue, we compared subcortical tissue (containing lateral ventricle-adjacent tissue) against superficial cortex, or hippocampus against superficial cortex; in rapid autopsy tissue, we compared lateral ventricle against superficial temporal cortex, or hippocampus against superficial temporal cortex **(Fig. 5a)**. The proportion of A2B5^+^EGFR^+^ cells was consistently enriched in the lateral ventricle and hippocampus compared to superficial cortex, in both young and aged donors **(Fig. 5b; Extended Data Fig. 10)**. Finally, the relative frequency of A2B5^+^EGFR^+^ cells across aging declined exponentially over the first two decades of life, but appeared to plateau afterwards **(Fig. 5c)**, which could represent a decline in the absolute number of NSCs with age, or a relative increase in the number of differentiated neural cells.

**Figure 5:**
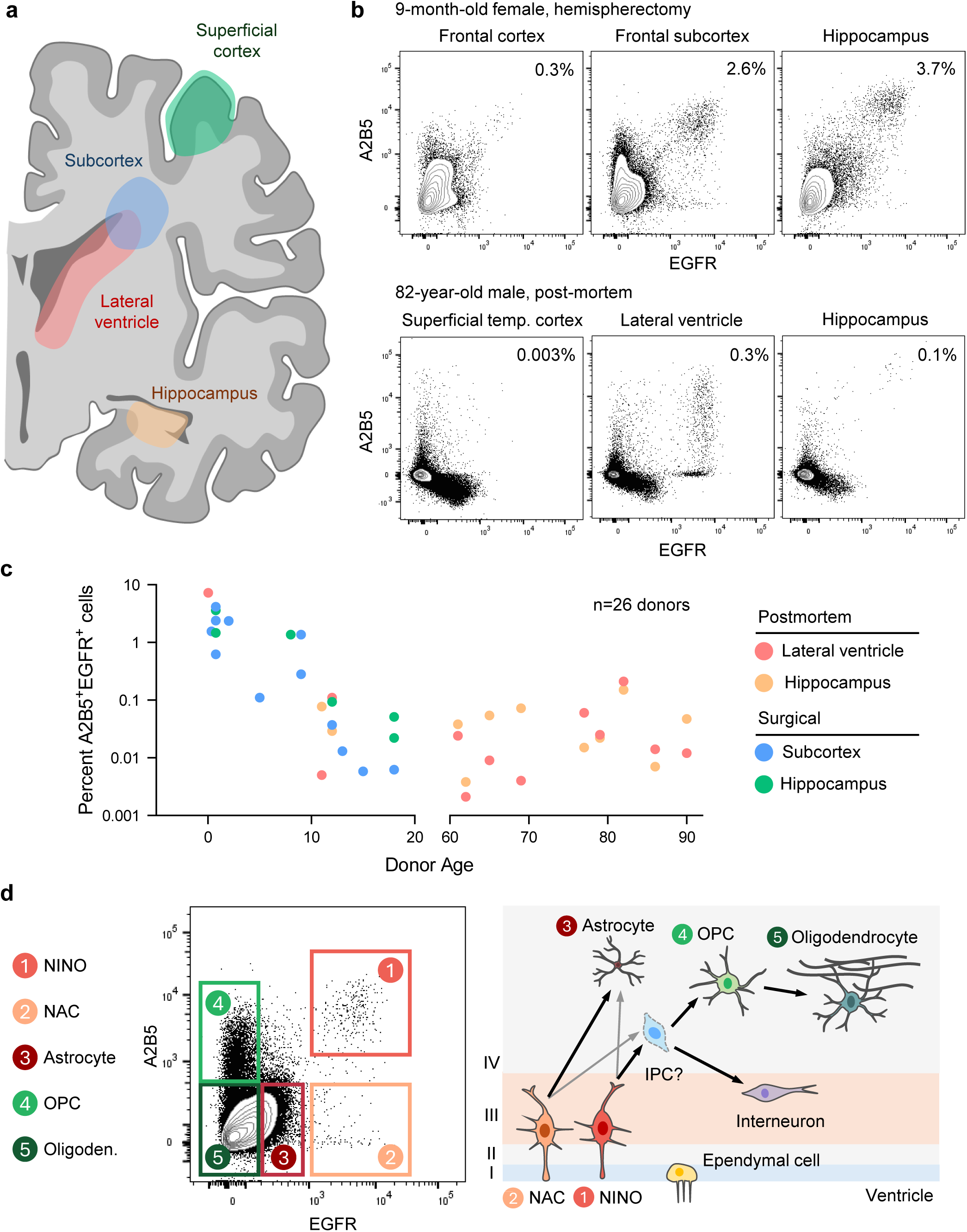
A2B5^+^EGFR^+^ putative NSCs are enriched in classically neurogenic zones. **a,** Schematic of coronal brain section showing representative regions from which tissue was obtained. **b,** Flow cytometry plots of dissociated brain tissue from a 9-month-old female and 82-year-old male, from various brain regions. Percentage of A2B5^+^EGFR^+^ events shown in top right corner. Events are pregated on live singlets of neural lineage. **c,** Frequency of A2B5^+^EGFR^+^ events in neurogenic zones, arranged by donor age. **d,** Schematic of proposed prospective isolation strategy and hierarchy of postnatal human NSCs.

## DISCUSSION

In this study, we identified and then purified bona fide neural stem cells from postnatal human brain tissue **(Fig. 5d)**. Single cell transcriptomics first identified a distinct population expressing genes suggestive of an NSC identity. Index sort analysis mapped these transcriptomically-defined NSCs onto their surface marker profile (immunophenotype), allowing for their isolation and experimental manipulation. The self-renewal and multilineage differentiation capability of these NSCs were demonstrated using both in vitro clonal differentiation assays, as well as in vivo xenotransplantation models, fulfilling the two essential definitions of stemness.

In the mouse, prospective isolation of NSCs from the SVZ is a well-established paradigm^49,50^, Most such protocols utilize transgenic lines, most commonly a GFAP::GFP reporter to mark both NSCs and astrocytes, combined with surface markers CD133 to mark NSCs, EGFR to mark activated NSCs, and CD24 to mark neuroblasts. Our study could not fully utilize these known markers, as papain-based tissue dissociation cleaves markers like CD24. We were, however, able to use A2B5 and EGFR in combination to distinguish oligodendrocytes (A2B5^−^EGFR^−^), OPCs (A2B5^+^EGFR^−^), astrocytes (A2B5^−^EGFR^mid^), and two populations of NSCs: the interneuron/OPC-biased NINOs (A2B5^+^EGFR^+^), and the astrocyte-biased NACs (A2B5^−^EGFR^hi^). A2B5 has previously been used as a marker for glial progenitors in the mouse and human brain^43,51,52^, but interestingly, a minor population of A2B5^+^ cells derived from human white matter has been reported to generate neurons in vitro^42^. Our study resolves this apparent discordance, by further resolving the heterogeneity within the A2B5^+^ population into an EGFR^−^ subset enriched for OPCs, and an EGFR^+^ population consisting of tripotent NSCs. Despite both NINO and NAC subsets expressing high surface levels of EGFR, they were not observed to be in cell cycle to any appreciable extent. Thus, EGFR may not distinguish activated versus quiescent NSCs in humans as it does in mice. Indeed, surface markers often vary significantly between analogous cell types of different species, such as CD34 on hematopoietic progenitors. The functional heterogeneity we identified among NSCs is not without precedence^53,54^. Mouse B1 cells in the V-SVZ are known to produce at least 10 subtypes of olfactory bulb interneurons^55,56^. Interestingly, the anatomical position of the B1 cell along the ventricular wall correlates with what interneuron subtype they produce, but this lineage bias appears to be cell-intrinsic, based on heterotopic grafting experiments^57^. The nature of NSC heterogeneity in humans likely differs considerably from that in the mouse, given differences in postnatal neurogenesis and gliogenesis. Further studies will require careful dissections of specific periventricular regions, combined with functional readouts.

Another question arises in the origin of postnatal human NSCs. In the mouse, radial glia produce pre-B1 cells mid-development, around embryonic day (E) 13.5-15.5^58,59^. These pre-B1 cells, after being specified, remain relatively quiescent until their reactivation postnatally. Dentate gyrus NSCs, on the other hand, become distinct from their embryonic radial glia precursors around the second or third postnatal week^60,61^. Human neurodevelopment has some notable innovations over that in mouse. In the second trimester, the ventricular radial glia (vRG) split into two pools: truncated radial glia (tRG), which lose their basal fiber and remain near the ventricular zone, and outer radial glia (oRG), which retain their basal fiber and situate in the outer subventricular zone (OSVZ)^2,62^. While it may be presumed that postnatal human NSCs similarly arise from radial glia, it is unknown at what time they are specified, and from what subset of radial glia. It is interesting that the NAC subset shares marker genes with oRG (*TNC*, *FAM107A*, *CLU*)^63^, but whether this observation reflects any developmental relation or merely coincidence could warrant further investigation. The window of human brain development from the third trimester to early postnatal life, and with it the transformation of the embryonic VZ into the adult V-SVZ, remains relatively elusive, in part due to scarcity of tissues. Further interrogation of this time period will prove critical to understanding the specification and transformation of prenatal to postnatal NSCs.

The identification of NSCs in the early postnatal human brain naturally raises the question of their fate later in life. While our data show that the frequency of A2B5^+^EGFR^+^ cells declines exponentially over the first two decades of life, it appears to stabilize thereafter, with the population identifiable even in a 90-year-old donor with dementia. This decline parallels that of B1 cells in the mouse V-SVZ, which deplete over lifespan due to consuming divisions^64,65^. On top of diminished numbers, aging is also associated with cell-intrinsic defects in NSC self-renewal, differentiation, and migration in mice^17,66–68^. How human NSCs change with aging, and whether they continue to generate neurons and glia to a physiologically relevant degree later in life, will certainly be of future scientific interest. Additionally, our isolation of postnatal NSCs will facilitate the study of their potential roles in pathology^24,25,69–71^, including psychiatric disorders, neurodegeneration, and glial neoplasms. Whether therapeutic targeting of NSCs is a feasible strategy for regenerative medicine would certainly be worthy of future exploration^72^.

## MATERIALS AND METHODS

### Mice

Work involving mice was carried out under the oversight of the Administrative Panel on Laboratory Animal Care (APLAC) at Stanford University, in compliance with approved protocols. Mice were housed at Stanford School of Medicine in Veterinary Service Center (VSC) facilities. Immunodeficient NSG (NOD.Cg-Prkdc^scid^ Il2rg^tm1Wjl^/SzJ, JAX: 005557) mice were used as recipients for intracranially transplanted human postnatal NSPCs. Mice were maintained in a 12:12 light/dark cycle at 18-23C, 40-60% humidity, with food and water given ad libitum. Mice were fed a rodent diet consisting of 18% protein / 6% fat (Envigo, cat. # 2918), with an irradiated high-fat diet consisting of 19% protein and 9% fat (Envigo, cat. # 2919) for pregnant and lactating dams. Neonatal mice (postnatal day 1-2) were transplanted with no restriction on sex. Litters were weaned between postnatal day 21-24 and housed with a maximum of 5 mice per cage, separated by sex.

### Human brain tissue

#### Collection

Fresh human brain tissue was collected shortly after death from deidentified donors. Postmortem tissue was obtained from two sources: (1) the Research Autopsy Center at Stanford (RACS), in accordance with Institutional Review Board (IRB) (IRB 63818); and (2) the International Institute for the Advancement of Medicine (IIAM), in coordination with the Indiana Donor Network and in compliance with research tissue collection protocols. Fresh surgical tissue was obtained from subjects enrolled in the Neuroscience Brain Bank (IRB #12625) undergoing medically indicated neurosurgical resection at Stanford Lucile Packard Children’s Hospital. For live-cell assays (culture and/or xenotransplantation), any donor with a history of central nervous system neoplasm was excluded. For RACS cases, formalin-fixed paraffin-embedded (FFPE) brain tissue was also provided.

#### Dissection

After recovery from the donor, postmortem brain samples were dissected by neuropathologists into the following regions: Lateral ventricle, hippocampus with subiculum, and superficial temporal lobe. In some instances, the dentate gyrus was removed and analyzed separately from the rest of the hippocampus. Where possible, surgical tissue resected en bloc was separated by anatomical region, including lobe of origin (eg frontal, temporal); deep (containing subventricular tissue) vs. superficial; and hippocampus if available. Anatomically dissected regions were placed into separate containers in hypothermasol media (STEMCELL Technologies, cat. # 07935) and maintained on ice.

### Adult brain tissue dissociation

All steps were performed using aseptic technique in a biosafety cabinet, with samples maintained on ice except for enzymatic digestion. Meningeal membranes, blood clots, and cauterized regions were removed using forceps and discarded. Brain tissue was weighed and transferred to a mincing buffer composed of calcium- and magnesium-supplemented Hank’s balanced salt solution (HBSS; Thermo Fisher Scientific, cat. # 24020117) combined with 0.1% polyvinyl alcohol (PVA) (Sigma, cat. # P8136), 10mM HEPES (Thermo Fisher Scientific, cat. # 15630080), 0.1% Poloxamer 188 (Thermo Fisher Scientific, cat. # 24040032), and 50μg/mL DNAse I (Worthington Biochemical, cat. # LS002007). Tissue was minced with razor blades in a vertical chopping motion to approximately 2mm^3^ pieces.

For enzymatic dissociation, tissue was digested in papain enzyme (Worthington Biochemical, cat. # LK003178) reconstituted in mincing buffer (see above) at a concentration of 20 units papain per ml. 4ml papain solution was added per 1g tissue. Tissue was digested at 37°C on a shaker for 15-25 min or until mixture passed smoothly several times through a P1000 pipet tip. Digested tissue was rinsed, pelleted, and resuspended in freshly diluted debris removal solution (Miltenyi Biotec, cat. # 130-109-398) prepared in cold Dulbecco’s phosphate-buffered saline (DPBS; Thermo Fisher Scientific, cat. # 14190250) according to the manufacturer’s handbook. 12ml diluted debris removal solution was used per gram of tissue, and transferred to 15ml tubes at a volume up to 8ml cell suspension per tube. 4ml cold DPBS was gently layered on top. Demyelination was performed by centrifuging tubes at 3,000 x g for 10 min (full acceleration and brake); aspirating top layer and interface containing myelin; resuspending cell pellets in cold DPBS; and centrifuging at 1,000 x g for 5 min. Supernatant containing any leftover debris removal solution was discarded.

To deplete red blood cells, tissue was resuspended directly in ACK lysing buffer (Thermo Fisher Scientific, cat. # A1049201) at approximately 2ml per gram of tissue, and incubated at room temperature for 2 min. The lysis reaction was halted by adding tenfold volume of 0.1% PVA HBSS buffer. Except where noted, all washes were performed in calcium- and magnesium-free HBSS (Thermo Fisher Scientific, cat. # 14175103) supplemented with 0.1% PVA. Dissociated brain tissue was incubated on ice in mincing buffer for 1-2 h to allow for antigen recovery, and rinsed/pelleted prior to antibody staining.

### Fluorescence-activated cell sorting (FACS)

For human samples, brain tissue was enzymatically dissociated into a single-cell suspension as described in the “Adult brain tissue dissociation” section. Cells were resuspended in human FcR blocking reagent (Miltenyi Biotec, cat. # 130-059-901) diluted 1:40 in 0.1% PVA HBSS buffer and incubated on ice for 5 min. Cell suspension was then stained on ice for 25 min with fluorophore-conjugated antibodies against A2B5 (clone 105HB29; Miltenyi Biotec, cat. # 130-123-800), EGFR (clone AY13; BioLegend, cat. # 352910), CD90 (clone 5E10; BioLegend, cat. # 328108), CD45 (clone HI30; BioLegend, cat. # 304010), CD11b (clone M1/70; BioLegend, cat. # 101242), CD31 (clone WM59; BioLegend, cat. # 303130), CD34 (clone 581; BioLegend, cat. # 343534), CD105 (clone 266; BD Biosciences, cat. # 562380), CD235a (clone HI264; BioLegend, cat. # 349120), and CD66b (clone G105F; BioLegend cat. # 305122), each used at a 1:50 dilution, except for anti-A2B5 which was used at 1:25. Cells were rinsed and resuspended in 0.1% PVA HBSS buffer. Propidium iodide (PI; Sigma, cat. # P4170) was added immediately prior to analysis at a final concentration of 1 µg/mL for exclusion of non-viable cells.

Flow cytometric analysis and cell sorting were performed on a FACSAria II instrument (BD Biosciences). Cellular debris was excluded based on forward- and side-scatter area parameters. Doublets were removed using a stringent two-step gating strategy based on forward- and side-scatter height versus width. For downstream scRNA-seq assays, cells were sorted using the highest purity (“single-cell”) mode. Bulk populations intended for cell culture and transplantation were collected using the “four-way purity” sort setting.

For mouse tissue, a FACS workflow was used to extract intracranially transplanted human cells from host brains for scRNA-seq. Fresh adult mouse brain tissue was single-cell dissociated as described in the “Adult brain tissue dissociation” section, and stained on ice for 25 min with fluorophore-conjugated antibodies against mouse CD45 (clone 30-F11; BioLegend, cat. # 103110), mouse CD11b (clone M1/70; Thermo Fisher Scientific, cat. # 15-0112-82), and TER-119 (clone TER-119; BioLegend, cat. # 116210), all at a 1:50 working dilution. Cells were rinsed, stained with PI, and gated by flow cytometry for live singlets as described above. Human cells were identified based on EGFP reporter expression and sorted on single-cell mode.

### Primary cell culture

#### NSC maintenance/expansion

Acutely isolated NSPCs were plated on Cultrex substrate (R&D Systems, cat. # 3434-010-02) in stem cell media composed of BrainPhys Neuronal Medium (STEMCELL Technologies, cat. # 05790) supplemented with 1% N-2 (Thermo Fisher Scientific, cat. # 17502048), 2% B-27 without vitamin A (Thermo Fisher Scientific, cat. # 12587010), 20ng/ml recombinant human epidermal growth factor (EGF; Fujifilm, cat. # PB-500-17), 20ng/ml recombinant human fibroblast growth factor 2 (FGF-2; Fujifilm, cat. # PB-500-17), 2μg/ml Heparin (STEMCELL Technologies, cat. #07980), and 63μg/ml N-acetylcysteine (VWR, cat. # E-3710). For enhanced in vitro expansion, stem cell medium was additionally supplemented with 20% Knockout serum replacement (Thermo Fisher Scientific, cat. # 10828010). Cultured NSPCs were dissociated with Accutase (Innovative Cell Technologies, cat. # AT104) for 6-8 min at room temperature and rinsed with 0.1% PVA HBSS buffer.

#### In vitro differentiation

For the evaluation of differentiation potential, primary NSPCs (either acutely isolated or after expansion in stem cell media) were plated on growth-factor-reduced Matrigel substrate (Corning, cat. # 356231). Three types of differentiation media were used: (1) growth-factor-free blank differentiation media consisting of a 1:1 mixture of DMEM/F12 (Thermo Fisher Scientific, cat. # 11320033) and Neurobasal-A (Thermo Fisher Scientific, cat. # 10888022) supplemented with 1% N-2 and 2% B-27; (2) neuronal medium additionally supplemented with 10ng/ml recombinant human brain-derived neurotrophic factor (BDNF; PeproTech, cat. # 450-02), 10ng/ml recombinant human glial cell line-derived neurotrophic factor (GDNF; PeproTech, cat. # 450-10), 10μM forskolin (Tocris, cat. # 1099), 10μM RO4929097 (Selleck Chemicals, cat. # S1575), and 200μg/ml 2-Phospho-L-ascorbic acid trisodium salt (AA2P; Sigma, cat. # 49752); and (3) glial media consisting of DMEM/F12 supplemented with 1% N-2, 2% B-27, 60ng/ml 3,3′,5-Triiodo-L-thyronine sodium salt (T3; Sigma, cat. # T6397), 10ng/ml recombinant human neurotrophin-3 (NT-3; PeproTech, cat. # 450-03), 10ng/ml recombinant human platelet-derived growth factor AA (PDGF-AA; Fujifilm, cat. # 100-16), 10ng/ml recombinant human insulin-like growth factor 1 (IGF-1; PeproTech, cat. # 100-11), 1μM dibutryl-cAMP (R&D Systems, cat. # 1141), 200μM L-ascorbic acid (Sigma, cat. # A92902), and 0.1% Trace Elements B Solution (Corning, cat. # 25022CI). Cells were treated with differentiation media for 5 days prior to cell fate analysis.

### Immunofluorescence

Cultured cells were fixed for 10 min at room temperature in 4% paraformaldehyde (PFA; Electron Microscopy Sciences, cat. # 15710) freshly prepared in PBS. Samples were gently washed 3 times with tris-buffered saline (TBS; Thermo Fisher Scientific, cat. # 28358) and then treated with a blocking/permeabilization buffer consisting of TBS supplemented with 0.3% Triton X-100 (Sigma, cat. # X100) and 3% horse serum (Corning, cat. # 35-030-CV) for 45 min at room temperature. Cells were incubated for 2 h at room temperature in a cocktail of primary antibodies diluted in stain buffer consisting of TBS supplemented with 0.3% Triton and 1% horse serum. Primary antibodies included: human GFAP (1:1000, Takara, clone STEM123, cat. #Y40420), OLIG2 (1:500, Abcam, clone EPR2673, cat. # ab109186), O4 (1:500, R&D Systems, cat. #MAB1326), SOX2 (1:1,000, Abcam, cat. # ab97959), and NESTIN (1:250, Thermo Fisher Scientific, clone SN06-27, cat. # MA5-32272). After 3 TBS washes, samples were incubated with the appropriate secondary antibodies diluted in stain buffer for 1.5 h at room temperature (Thermo Fisher Scientific). Samples were stained for 10 min at room temperature with DAPI (Thermo Fisher Scientific, cat. # D1306) at a concentration of 1 mg/mL in PBS, and rinsed 3 times in TBS. Images were captured on a Leica DMi8 inverted microscope or a Molecular Devices ImageXpress Micro 4 high-content imaging system. For O4 staining, cells were exposed to primary antibody for 30 min at 37°C prior to fixation.

### Lentiviral Barcoding

STICR lentivirus was transfected using a second-generation lentiviral packaging system comprising pMD2.G (Addgene #12259) and psPAX2 (Addgene #12260). To enhance viral titer, a pcDNA3.1 puro Nodamura B2 plasmid (Addgene #17228) was co-transfected with the packaging and transfer plasmids. Plasmids were introduced into HEK-293 cells (ATCC, cat. # CRL-1573) using TransIT-293 transfection reagent (Mirius Bio, cat. # MIR 2700). HEK-293 cells were transfected in serum-free conditions using Opti-MEM (Thermo Fisher Scientific, cat. # 31985070) without any supplements.

Prior to transfection, HEK-293 cells were maintained in high-glucose DMEM (Thermo Fisher Scientific, cat. # 10566024) supplemented with 10% fetal bovine serum (HyClone, cat. #SH30071.03HI) and 1% penicillin–streptomycin (Thermo Fisher Scientific, cat. # 15070063). They were split after achieving 50-70% confluency using 0.25% Trypsin-EDTA (Thermo Fisher Scientific, cat. # 25200056) for 3-5 min at 37C.

Supernatant containing lentiviral particles was harvested 24 h and again at 48 h post-transfection, filtered through a 0.45 µm filter (Millipore Sigma, cat. # SE1M003M00), and combined at a 4:1 ratio with 50% polyethylene glycol solution (PEG; need catalog number) prepared in HBSS. Viruses were precipitated in PEG for 24 h at 4C and then pelleted by centrifugation at 2,500 x g for 20 min. Viral pellets were resuspended in 250 µL HBSS, aliquoted, and stored at −80 °C. Human NSPCs were virally transduced for 1.5 h at 37C on a shaker.

### Xenotransplantation

After brief (∼2 w) expansion in vitro, primary human NSPCs were resuspended in HBSS supplemented with 0.1% Fast Green dye (Sigma, cat. # F7252) to facilitate visualization of the injection site. Neonatal NSG mice (postnatal day 1–2) were randomized, cryoanesthetized on ice for 2 min, and positioned in a stereotaxic apparatus (Harvard Apparatus) equipped with a neonatal mouse adaptor (Cunningham). The sinus above lambda was identified under transillumination and used as the anatomical reference point. Burr holes were created in the skull cartilage at the target sites using a 30-gauge needle. Cell suspensions were delivered bilaterally into the lateral ventricles using a 10μl calibrated syringe (Hamilton, cat. # CAL7653-01) fitted with a 33-gauge needle (Hamilton, cat. # 7803-15) and controlled by a Micro4 microsyringe pump (World Precision Instruments) at an infusion rate of 1 µL/min, with 1.5 µL injected per ventricle. Accurate ventricular delivery was confirmed visually using light illumination. Stereotaxic coordinates relative to lambda were as follows: anteroposterior from midline (A), lateral from midline (L), ventral from brain surface (V). Lateral ventricles (A, L, V) = (0.8, ±1.5, 2.0) mm with reference to lamda. Human-to-mouse xenograft experiments with primary postnatal human brain cells were carried out with approval from the Stem Cell Research Oversight (SCRO) at Stanford University, in compliance with SCRO protocol #735.

### Histology

For xenografted mouse brains, mice were sacrificed 6 months after transplantation and perfused with HBSS containing 5mM EDTA (Thermo Fisher Scientific, cat. # 15575020). The brain was removed, cut in half down the midline, and placed into 4% paraformaldehyde (Electron Microscopy Sciences, cat. # 15710) freshly prepared in PBS. Brains were fixed for 16-20 h at 4C, rinsed in PBS, and transferred to 30% sucrose solution (Fisher Scientific, cat. # BP220-212) for at least 24h at 4C. Brains were cut into 40 μm sagittal sections using a sliding microtome (Leica SM2010R). Tissue slices were stored prior to staining at −20°C in tissue collecting solution consisting of 25% glycerin (vol/vol), 30% ethylene glycol, 1.38g/L NaH_2_PO_4_, and 5.48g/L Na_2_HPO_4_. Floating brain sections were stained using the “immunofluorescence” protocol, described above, with the following modifications: (1) samples were exposed to primary antibodies overnight at 4°C, with gentle agitation; and (2) tissue sections were transferred to 0.1 mM phosphate buffer (Millipore Sigma, cat. # P3619) after the final TBS rinse, manually arranged on microscope slides, air-dried, and coverslip-sealed in ProLong Gold anti-fade mountant (Thermo Fisher Scientific, cat. # P36934). In addition to those listed in the “Immunofluorescence” section, antibodies used in mouse xenograft slices included human cytoplasmic antigen STEM121 (1:500, Takara, cat. # Y40410), NeuN (1:500, Abcam, cat. # ab104225), OLIG2 (1:350, R&D Systems, cat. # AF2418), human EGFR (1:100, R&D Systems, cat. # AF231), GFAP (1:500, Abcam, cat. # ab7260), and MAP2 (1:1,000, Abcam, cat. # ab5392).

For primary human brain samples, 5 μm sections of human FFPE brain tissue were stained either by a single immunohistochemistry (IHC) workflow or a dual-labeling protocol combining both IHC and RNA in-situ hybridization (ISH). Formalin-fixed paraffin-embedded (FFPE) human donor samples were obtained from the Stanford Department of Pathology and sectioned at 5 µm by HISTO-TEC LABORATORY INC prior to in situ hybridization experiments. RNAscope in situ hybridization was performed using the RNAscope Multiplex Fluorescent v2 assay (Advanced Cell Diagnostics/Bio-Techne, Newark, CA.) according to the manufacturer’s instructions, with minor modifications as described below. Slides were baked for 1 h at 60 °C and deparaffinized by two washes in xylene (Sigma-Aldrich, cat. # 534056) followed by two washes in 99.5% ethanol (Thermo Scientific Chemicals, cat. # 615090040). Deparaffinized slides were incubated with RNAscope Hydrogen Peroxide (Advanced Cell Diagnostics/Bio-Techne, cat. # 323270) and then boiled in RNAscope 1× Target Retrieval Reagent (Advanced Cell Diagnostics/Bio-Techne, cat. # 323166) for 20 min.

After target retrieval, a hydrophobic barrier was drawn around the tissue using an Immedge hydrophobic barrier pen (Vector Laboratories, cat. # H-4000). For combined immunolabeling, sections were blocked and incubated overnight at 4 °C with primary antibodies diluted 1:200 in a staining buffer (TBS supplemented with 0.3% Triton X-100 and 1% horse serum). Sections were then processed the following day according to the RNAscope Multiplex Fluorescent v2 workflow using a probe against human *EGFR* mRNA in channel 1 (ACD, cat. # 310061) with hybridization performed at 40 °C in a HybEZ oven. Signal amplification and detection were carried out following the manufacturer’s protocol using TSA Vivid Fluorophore 650 (ACD, cat. # 323110).

Finally, sections were incubated at room temperature for 1 h 20 min with secondary antibodies diluted 1:500 in staining buffer. Nuclei were counterstained with DAPI diluted 1:1000 in 1× TBS for 10 min before coverslips were mounted onto the sample using ProLong Gold anti-fade mountant.

### Smart-seq3

#### Single-cell isolation and lysis

Brain tissue was processed into a single-cell suspension and prepared for FACS as described in the sections “Adult brain tissue dissociation” and “Fluorescence-activated cell sorting (FACS)”. Individual cells were index-sorted into 96-well plates preloaded with 2 µL of lysis buffer containing 1 U/µL RNase inhibitor (Clontech, cat. #2313B), 0.1% Triton X-100 (Thermo Fisher Scientific, cat. #85111), 2.5 mM dNTPs (Thermo Fisher Scientific, cat. #10297018), 2.5 µM oligo(dT)_30_VN primer (Integrated DNA Technologies; 5′-AAGCAGTGGTATCAACGCAGAGTACT_30_VN-3′), and ERCC RNA spike-in controls (Thermo Fisher Scientific, cat. #4456739) diluted 1:600,000 in UltraPure water (Thermo Fisher Scientific, cat. #10977015). Immediately following sorting, plates were centrifuged at 3,000 × g for 30 s at 4 °C, snap-frozen on dry ice, and stored at −80 °C until further processing.

#### Reverse transcription and cDNA pre-amplification

Reverse transcription (RT) and cDNA pre-amplification were carried out using a modified Smart-seq3 protocol [cite Hagemann-Jensen et al. Nat Biotech]. Lysis plates were thawed on ice, incubated at 72 °C for 3 min, and immediately placed on ice to facilitate oligo(dT)_30_VN primer annealing. Reverse transcription was initiated by adding 3 µL of RT reaction mix to each well using a Mantis liquid handler (Formulatrix). The RT mix consisted of 25 mM Tris-HCl (pH 8.5; Teknova, cat. #T5085), 0.5 U/µL RNase inhibitor, 8 mM dithiothreitol (Promega, cat. #P1171), 30 mM NaCl (Thermo Fisher Scientific, cat. #AM9760G), 2.5 mM MgCl_2_ (Thermo Fisher Scientific, cat. #AM9530G), 1 mM GTP (Thermo Fisher Scientific, cat. #R0461), 2 µM template-switching oligonucleotide (TSO) (Integrated DNA Technologies; 5′-AAGCAGTGGTATCAACGCAGAGTGAATrGrGrG-3′), 5% polyethylene glycol (Sigma, cat. #P1458), and 2 U/µL Maxima H-minus reverse transcriptase (Thermo Fisher Scientific, cat. #EP0753) in UltraPure water. Reactions were incubated in a C1000 Touch thermal cycler (Bio-Rad) at 42 °C for 90 min, followed by enzyme inactivation at 70 °C for 15 min.

For pre-amplification, 7.5 µL of PCR mix containing 1.67× KAPA HiFi HotStart ReadyMix (Kapa Biosystems, cat. #KK2602) and 0.17 µM ISPCR primer (Integrated DNA Technologies; 5′-AAGCAGTGGTATCAACGCAGAGT-3′) was added to each well. Amplification was performed under the following cycling conditions: 98 °C for 3 min; 24 cycles of 98 °C for 20 s, 67 °C for 15 s, and 72 °C for 6 min; followed by a final extension at 72 °C for 5 min. Preamplified cDNA was purified using 0.65–0.75× volumes of AMPure XP beads (Beckman Coulter, cat. #A63882) to remove residual primers and fragments smaller than 400 bp, and eluted in 12.5 µL UltraPure water.

#### cDNA quality assessment and normalization

1 µL of purified cDNA from each well was transferred to a separate 96-well plate for quality control. cDNA concentration and fragment size distributions were assessed using a capillary electrophoresis–based Fragment Analyzer (Advanced Analytical). Wells with cDNA concentrations below 1.7 ng/µL were excluded; this threshold was established based on measurements from blank wells containing ERCC spike-ins without sorted cells. Wells exceeding the concentration cutoff were transferred to a new 384-well plate using a Mosquito X1 liquid handler (SPT Labtech). During reformatting, samples were diluted with UltraPure water to achieve normalized concentrations between 1.7 and 4.0 ng/µL.

#### Library construction and sequencing

Normalized cDNA was used as input for Illumina-compatible library preparation. Tagmentation reactions were assembled by combining 0.4 µL cDNA with 1.2 µL of a Tn5 transposase mix containing 1 ng/µL Tn5 enzyme, 16 mM Tris-HCl (pH 7.6), 16 mM MgCl_2_, and 8% dimethylformamide (Thermo Fisher Scientific, cat. #AC327171000) in UltraPure water. Reactions were quenched by the addition of 0.4 µL neutralization buffer containing 0.1% SDS.

Indexing PCR was performed by adding 0.4 µL each of 5 µM i5 and i7 indexing primers (Integrated DNA Technologies; custom 7,680-plex unique dual-index primer set) and 1.2 µL KAPA HiFi HotStart ReadyMix. Amplification was carried out in a C1000 Touch thermal cycler using the following program: 72 °C for 3 min; 95 °C for 30 s; 10 cycles of 98 °C for 10 s, 67 °C for 30 s, and 72 °C for 60 s. For each 384-well plate, 1 µL was taken from each well and combined. Pooled libraries were purified using 0.8× volume of AMPure XP beads. Library concentration and fragment size distributions were assessed using a Fragment Analyzer. Twenty 384-cell library pools were normalized for concentration, combined to generate a 7,680-plex library pool, and subjected to an additional cleanup and concentration step using 0.8× AMPure beads. Final libraries were sequenced on an Illumina NovaSeq 6000 using S4 flow cells to generate approximately 1–2 million paired-end 2 × 150 bp reads per cell.

#### Read alignment and quantification

Base call files were demultiplexed using bcl2fastq (v2.19.0.316). Adapter trimming of 3′ sequences was performed with Skewer (v0.2.2), after which reads were aligned to the human reference genome hg38 (GENCODE GRCh38.p13) using the STAR aligner (v2.6.1d) in two-pass mode. During the first mapping pass, reads from each cell were aligned against a STAR genome index constructed from GENCODE human transcript annotations (v34). Splice junctions identified during this initial alignment were collected across all cells and used to generate a revised STAR genome index that incorporated both annotated and newly detected splice junctions. This updated index was subsequently applied for second-pass read alignment.

STAR alignment parameters were based on recommendations from the ENCODE long-mRNA pipeline and the STAR documentation. In addition to ENCODE-suggested options, the “--quantMode TranscriptomeSAM” option was enabled during second-pass alignment to generate BAM files containing transcriptome-aligned reads. These BAM files served as input for expression quantification at both the gene level (aggregating expression across all annotated transcript isoforms) and the transcript level using RSEM (v1.3.3) with the “--single-cell-prior” option, which accounts for the sparsity characteristic of single-cell RNA-seq data.

#### Data preprocessing and downstream analysis

Gene-level count matrices were integrated with associated metadata using the Scanpy Python package (v1.8.2). Genes detected in fewer than three cells were excluded, as were cells with fewer than 500 expressed genes or fewer than 5,000 total read counts. Counts were normalized using size-factor normalization to a total of 10,000 reads per cell, log-transformed, and scaled with a maximum value capped at 10. Highly variable genes were identified using default Scanpy settings.

Dimensionality reduction was performed using principal component analysis, followed by construction of a nearest-neighbor graph and clustering with the Leiden algorithm. Pseudotime analysis was used to infer gene expression dynamics along putative maturation trajectories.

### PIP-seq

For cells carrying STICR lineage tracing constructs, single-cell transcriptomes and corresponding clonal barcodes were simultaneously profiled using Pip-seq. Manufacturer-provided instructions (Fluent Biosciences, doc. ID # FB0002130) were followed to generate single-cell gene expression libraries. For construction of STICR barcode libraries, 10 µL of PIP-seq full-length cDNA was used as input for a 50 µL PCR reaction containing 25 µL Q5 Hot Start High-Fidelity 2× Master Mix (NEB, cat. #M0494) and STICR barcode read 1 and read 2 primers (0.5 µM each), as previously described by Delgado *et al.* PCR amplification was carried out using the following thermal cycling conditions: initial denaturation at 98 °C for 30 s; 15 cycles of 98 °C for 10 s, 62 °C for 20 s, and 72 °C for 10 s; followed by a final extension at 72 °C for 2 min and a hold at 4 °C.

Following amplification, libraries underwent dual-sided size selection (0.6–0.8×) using SPRIselect beads (Beckman Coulter, cat. #B23318). Final libraries were sequenced on Illumina NovaSeq instruments to an average depth of approximately 25,000 reads per cell for gene expression libraries and 5,000 reads per cell for STICR barcode libraries.

#### Joint transcriptome and barcode alignment

Raw FASTQ files from cDNA and STICR libraries were processed with PIPseeker (v3.0) for read trimming and barcode extraction. Trimmed reads and barcode whitelists generated by PIPseeker were aligned using STARsolo (v2.7.11a) against optimized human and mouse reference genomes obtained from The Pool Lab (https://www.thepoollab.org/resources). STARsolo was executed in a configuration to enable barcode-aware UMI processing, empty-droplet filtering, probabilistic handling of multimapping reads, and generation of gene- and splicing-resolved outputs for downstream expression, velocity, and lineage analyses. Transcript count matrices were decontaminated for ambient RNA using the decontX workflow, with non-called cells identified by PIPseeker used to model the ambient RNA background.

#### STICR barcode processing and clonal assignment

STARsolo output BAM files were processed using a forked implementation of NextClone (https://github.com/cnk113/NextClone), which enables bitwise barcode whitelisting and filtering based on barcode UMI read coverage. Barcodes were retained if supported by at least 10 reads across the full sequencing library, and UMIs were quantified only if supported by a minimum of two reads. NextClone output was subsequently processed using CloneDetective to perform additional read-level UMI filtering.

Barcodes detected in ≥50% of called cells were blacklisted to exclude ambient barcode contamination. Residual ambient barcode signal was further removed using decontX applied to the barcode UMI–by–cell matrix. Cells were retained for clonal analysis if they expressed more than one barcode UMI. Clones containing fewer than three cells were filtered out. Clonal group assignment was performed using a hypergeometric test to assess barcode overlap, particularly in conditions with higher infection rates.

## ACKNOWLEDGMENTS

We deeply thank the patients and families who donated tissue for this study. We thank Aaron McCarty for mouse colony management; Teja Naik and Linda Quinn for laboratory management; Catherine Carswell-Crumpton, Cheng Pan, and Joe Pasillas for flow cytometry support; Katia Alvarez, Michael R. Eckart, and the Stanford Protein and Nucleic Acid (PAN) Facility for technical support with cDNA quality control; Honey Mekonen, Angela Detweiler, and the Chan Zuckerberg Biohub for sequencing support.

D.D.L. is supported by Stanford University Medical Scientist Training Program grant T32-GM007365 and T32-GM145402, and the Seth A. Ritch Bio-X Stanford Interdisciplinary Graduate Fellowship (SIGF). A.E.E. is supported by the ISCBRM Siebel Scholar Program. This study was carried out as part of Team NexTGen within the Cancer Grand Challenges initiative funded by Cancer Research UK (CGCATF-2021/100005), the National Cancer Institute (OT2CA278686), and The Mark Foundation for Cancer Research. This work was supported by an NIH/NCI Outstanding Investigator Award R35-CA220434 to I.L.W., a Stinehart-Reed Award to I.L.W., the Virginia and D.K. Ludwig Fund for Cancer Research to I.L.W. and to the Research Autopsy Center at Stanford (RACS), NIH R01-NS123263 to T.J.N., CIRM DISC0-14429 to T.J.N., Sontag Foundation Distinguished Scientist Award to T.J.N., and gifts from Schmidt Futures and the William K. Bowes Jr. Foundation to T.J.N. T.J.N is a NYSCF Robertson Neuroscience Investigator. L.M.P. is a Chambers-Okamura Faculty Scholar in Pediatric Neurosurgery.

## DATA AND CODE AVAILABILITY

The raw data FASTQ files and gene count table have been deposited in Synapse.

---

**Extended Data Fig. 1:**
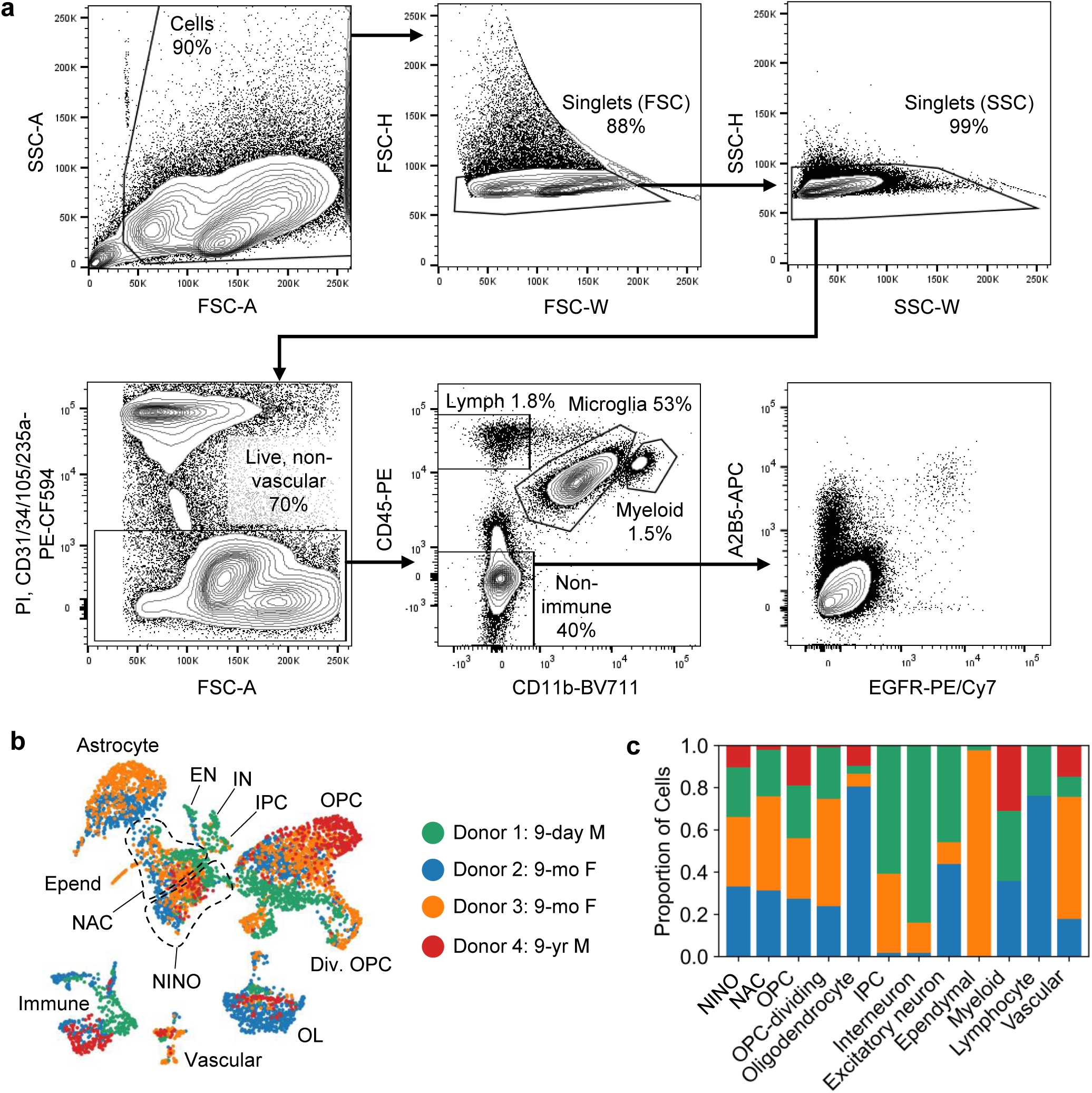
Flow cytometric gating of postnatal human brain cells. **a,** Flow cytometry plots showing pre-gating of dissociated postnatal human brain tissue, to exclude debris, doublets, dead cells, and non-neural lineages. **b,** UMAP plot of postnatal human brain cells, colored by donor. **c,** Stacked bar plot showing contribution of each donor to each transcriptomic cell cluster.

**Extended Data Fig. 2:**
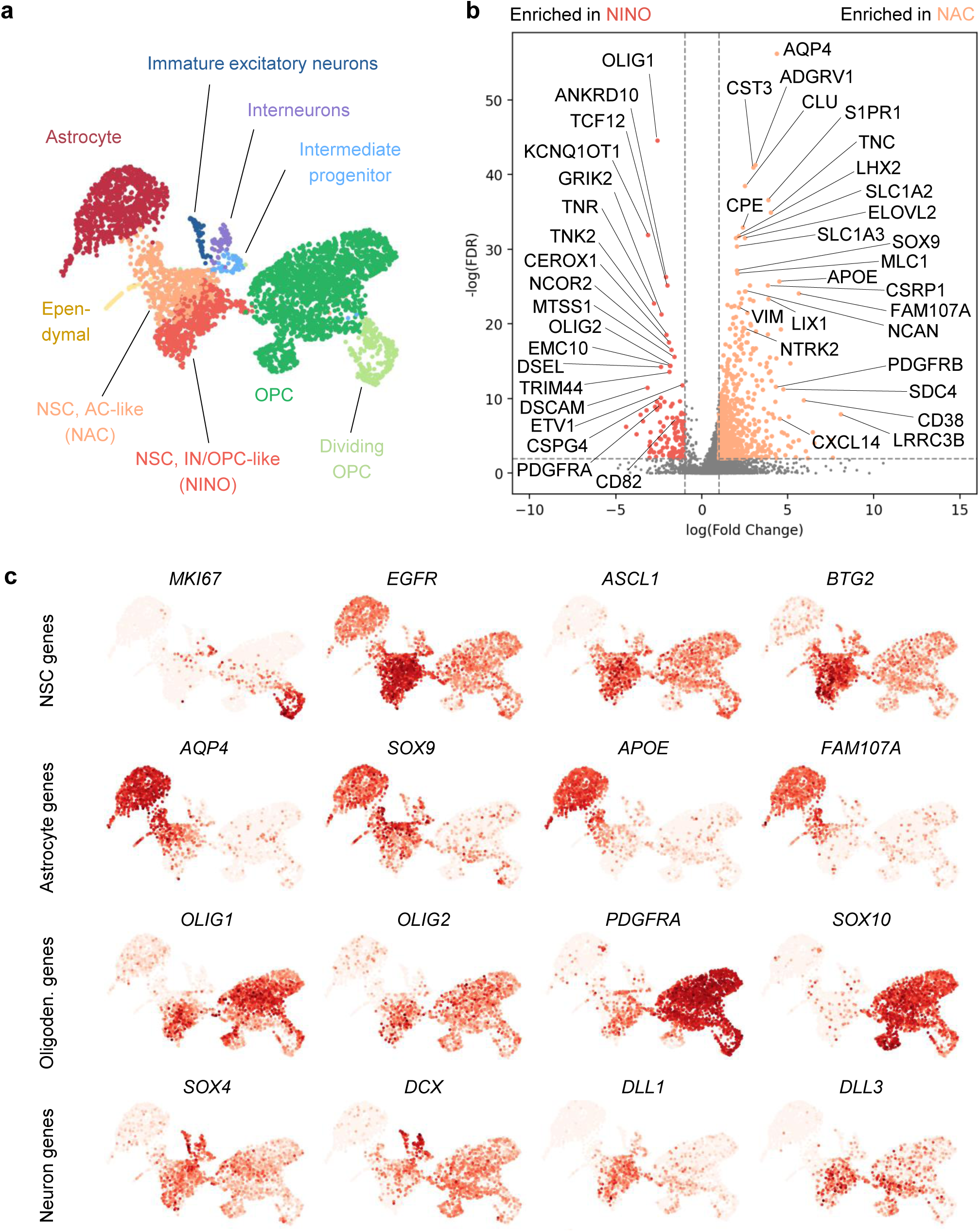
Transcriptomic heterogeneity among postnatal NSCs. **a,** UMAP plot with cluster annotations of postnatal human brain cells sequenced using Smart-seq3, subset on stem and progenitor clusters. **b,** Volcano plot sowing differentially-expressed genes between NINO and NAC clusters. **c,** UMAP plot with gene expression of selected marker genes.

**Extended Data Fig. 3:**
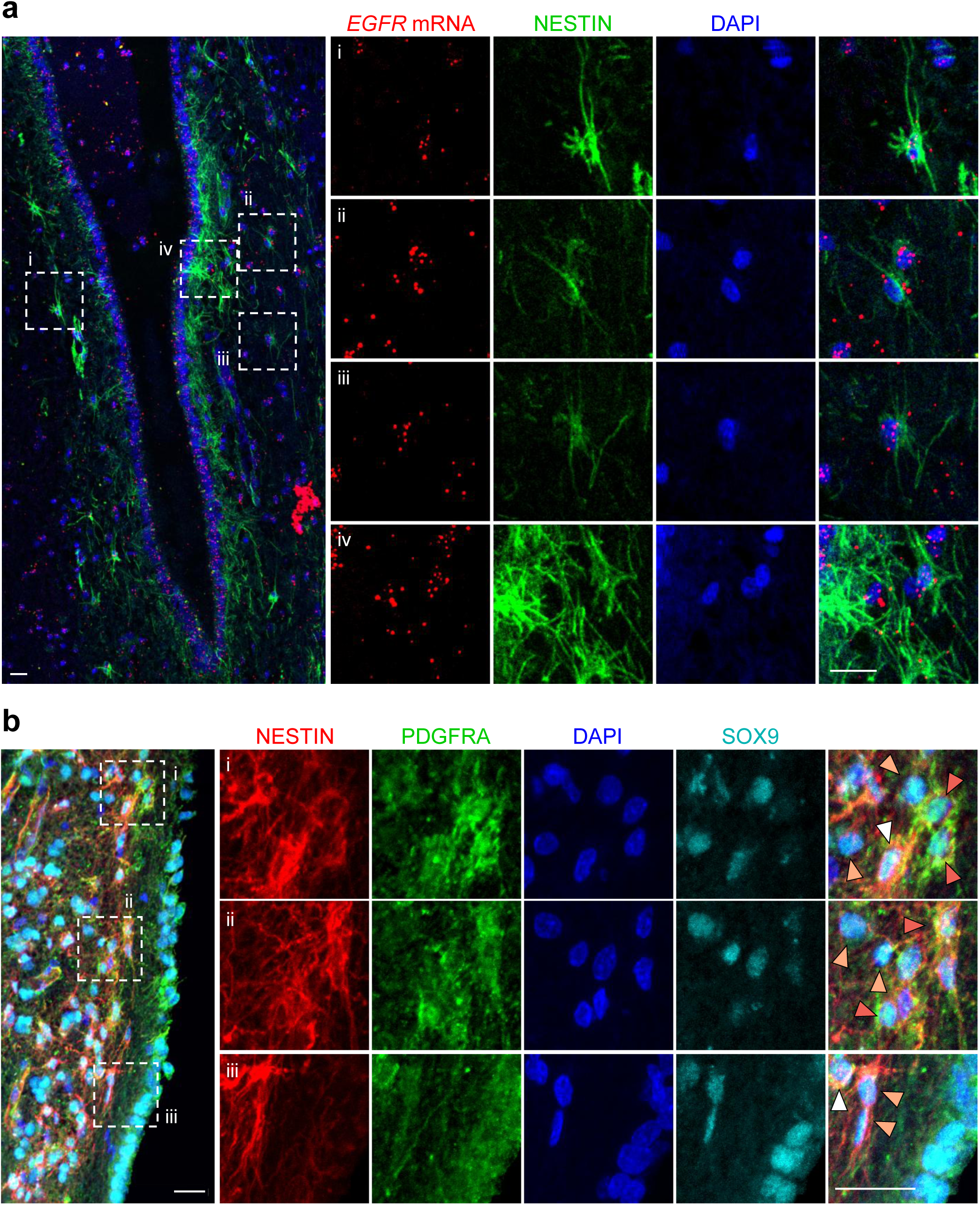
Histology of postnatal human NSCs in the SVZ. **a,** Human SVZ tissue from a 16-year-old donor, stained by immunofluorescence and in situ hybridization for DAPI (blue), *EGFR* mRNA (red), and NESTIN (green). **b,** Human SVZ tissue from a 65-year-old donor, stained by immunofluorescence for DAPI (blue), NESTIN (red), PDGFRA (green), and SOX9 (cyan). Arrowheads: NES^+^PDGFRA^+^SOX9^−^ (red); NES^+^PDGFRA^−^SOX9^+^ (orange); NES^+^PDGFRA^+^SOX9^+^ (white). Scale bars: 20 μm

**Extended Data Fig. 4:**
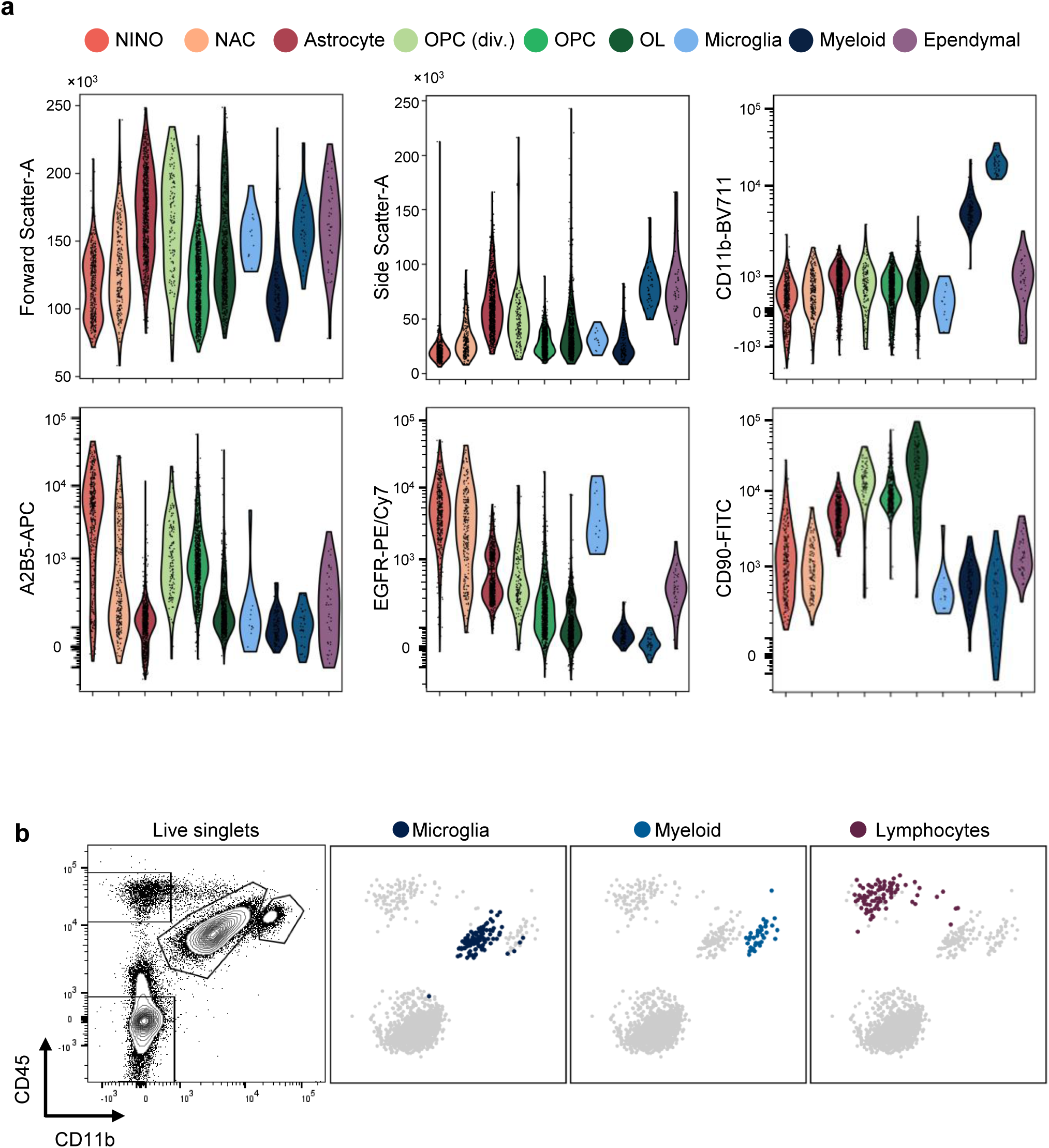
Index sort analysis of postnatal human brain cells. **a,** Violin plots showing mean fluorescence intensity (MFI) of surface markers measured by index sort, plotted per transcriptomic cell cluster. **b,** Index sort analysis mapping immune cell types onto their surface marker profile. Flow cytometry plot is pre-gated on live singlets.

**Extended Data Fig. 5:**
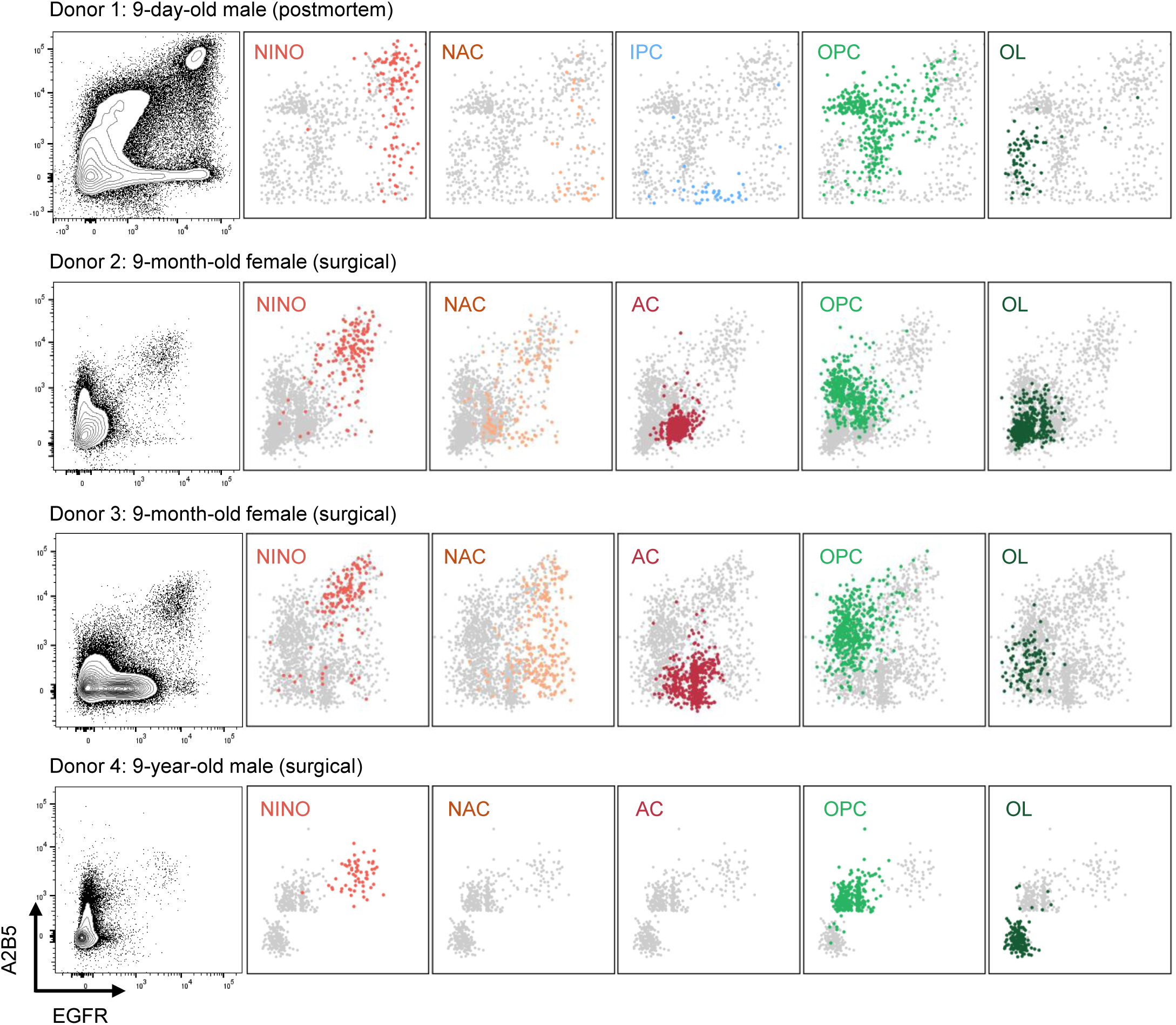
Index sort analysis of postnatal human brain tissue by donor. Index sort analysis mapping transcriptomically-defined clusters onto their surface marker profile (immunophenotype), separated by donor.

**Extended Data Fig. 6:**
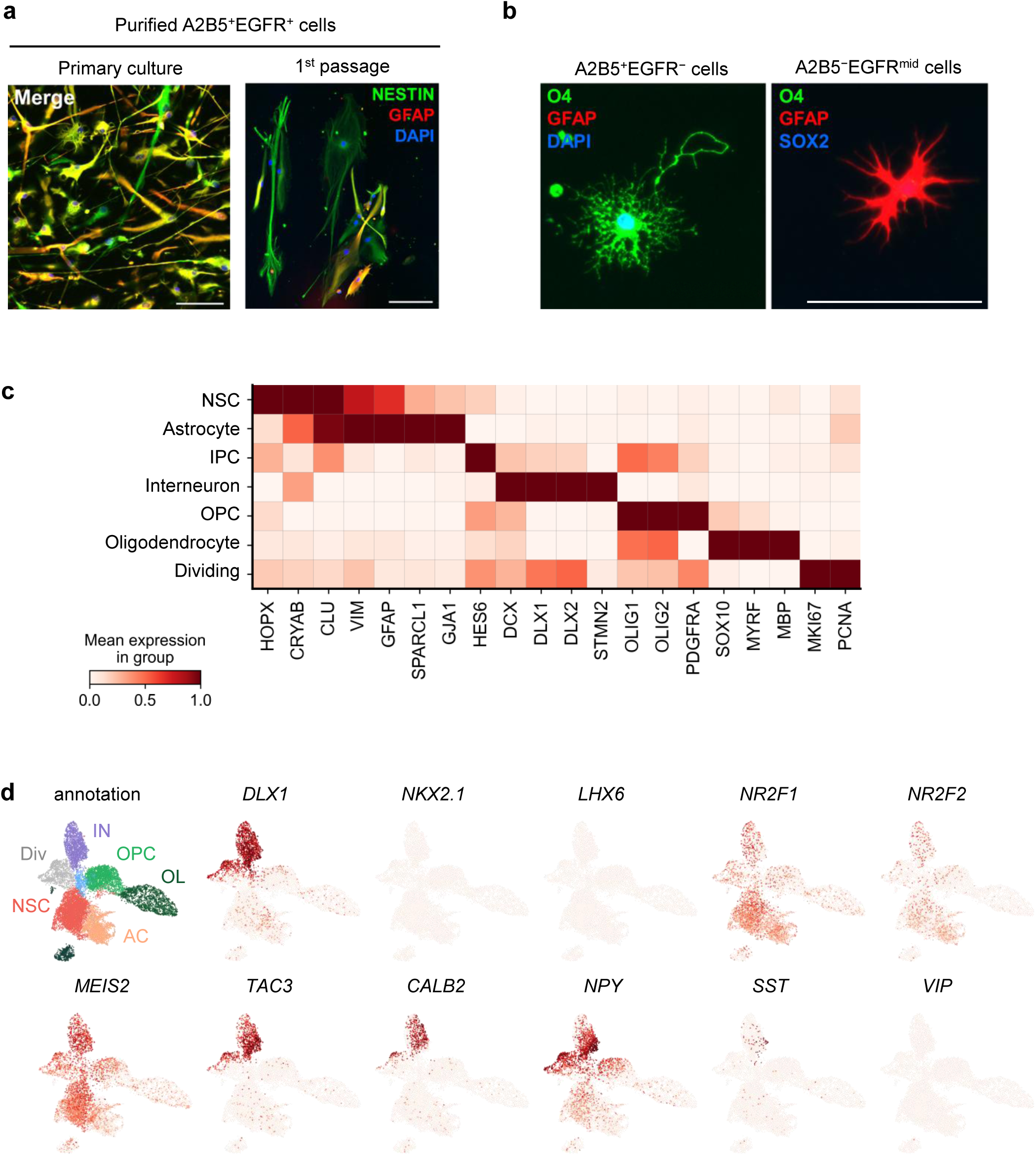
Profiling of cultured postnatal human NSCs. **a,** Immunofluorescence images of purified A2B5^+^EGFR^+^ cells at passage 0 (left) or following one passage (right). **b,** Immunofluorescence images of purified cells from the postnatal human brain at passage 0. Left, A2B5^+^EGFR^−^ cells stained for DAPI (blue), O4 (green), and GFAP (red). Right, A2B5^−^EGFR^mid^ stained for O4 (green), GFAP (red), and SOX2 (blue). Cells in **a** and **b** were derived from a 9-year-old male donor. Scale bars 100 μm. **c,** Matrix plot showing expression of marker genes by transcriptomic cluster from cultured human brain cells. **d,** UMAP plot with gene expression of selected interneuron marker genes.

**Extended Data Fig. 7:**
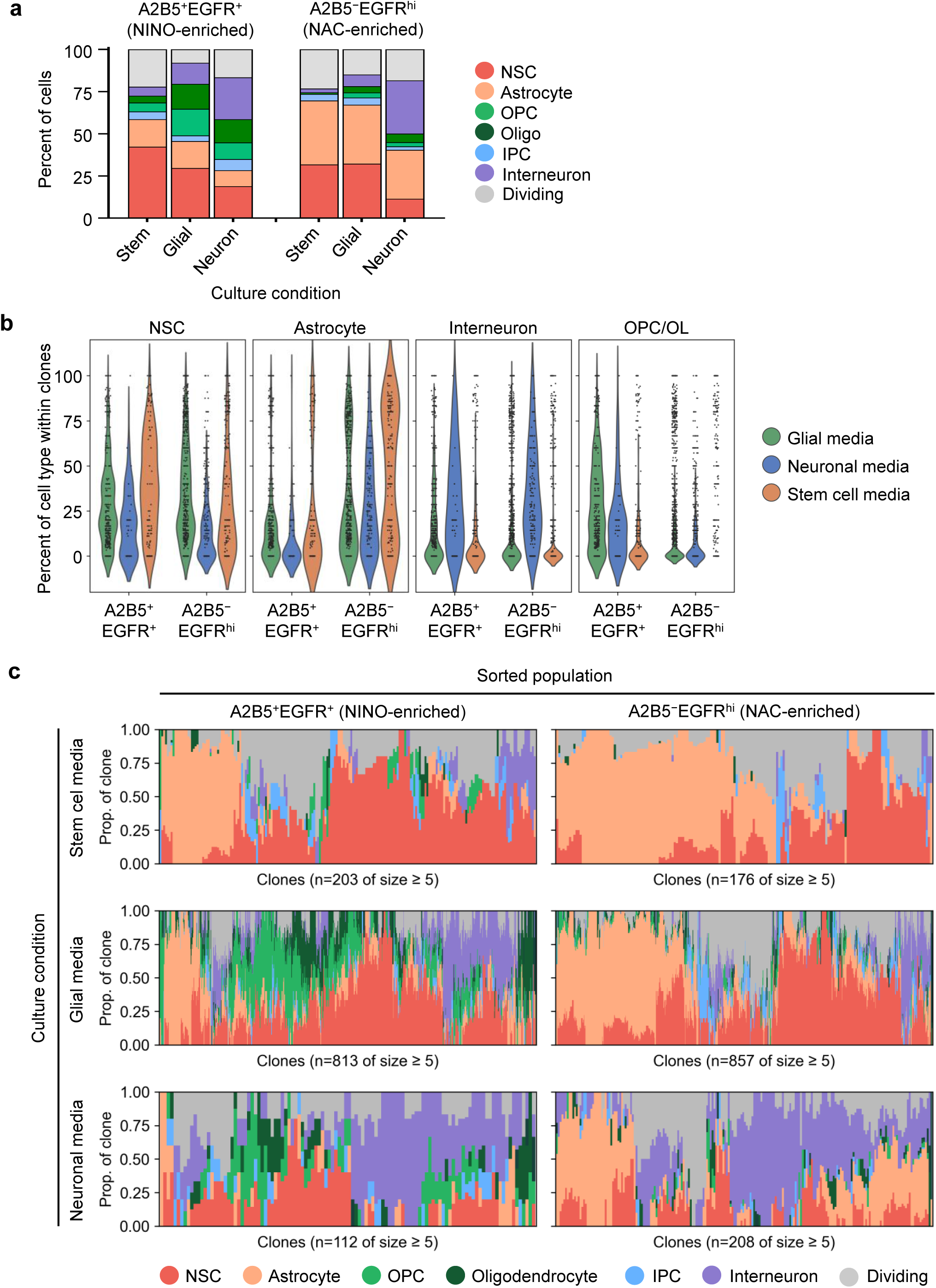
Postnatal human NSCs respond to differentiation signals. **a,** Stacked bar plot showing relative proportions of cell types resulting from in vitro culture of A2B5^+^EGFR^+^ or A2B5^−^EGFR^hi^ cells, in stem cell media, glial media, or neuronal media. **b,** Violin plots showing makeup of clones derived from cultured A2B5^+^EGFR^+^ or A2B5^−^EGFR^hi^ cells, in either stem cell media, glial media, or neuronal media. Each dot represents 1 clone of size ≥ 5. Position along y-axis represents what percent of cells within that clone are of the given cell type (NSC, astrocyte, interneuron, or OPC/oligodendrocyte). **c,** Stacked bar plots showing relative cell type makeup within each clone (size ≥ 5) recovered following culture. Within a plot, each column represents one clone. Plots are separated by sorted population (left, A2B5^+^EGFR^+^; right, A2B5^−^EGFR^hi^) and culture media (top, stem cell media; middle, glial media; bottom, neuronal media).

**Extended Data Fig. 8:**
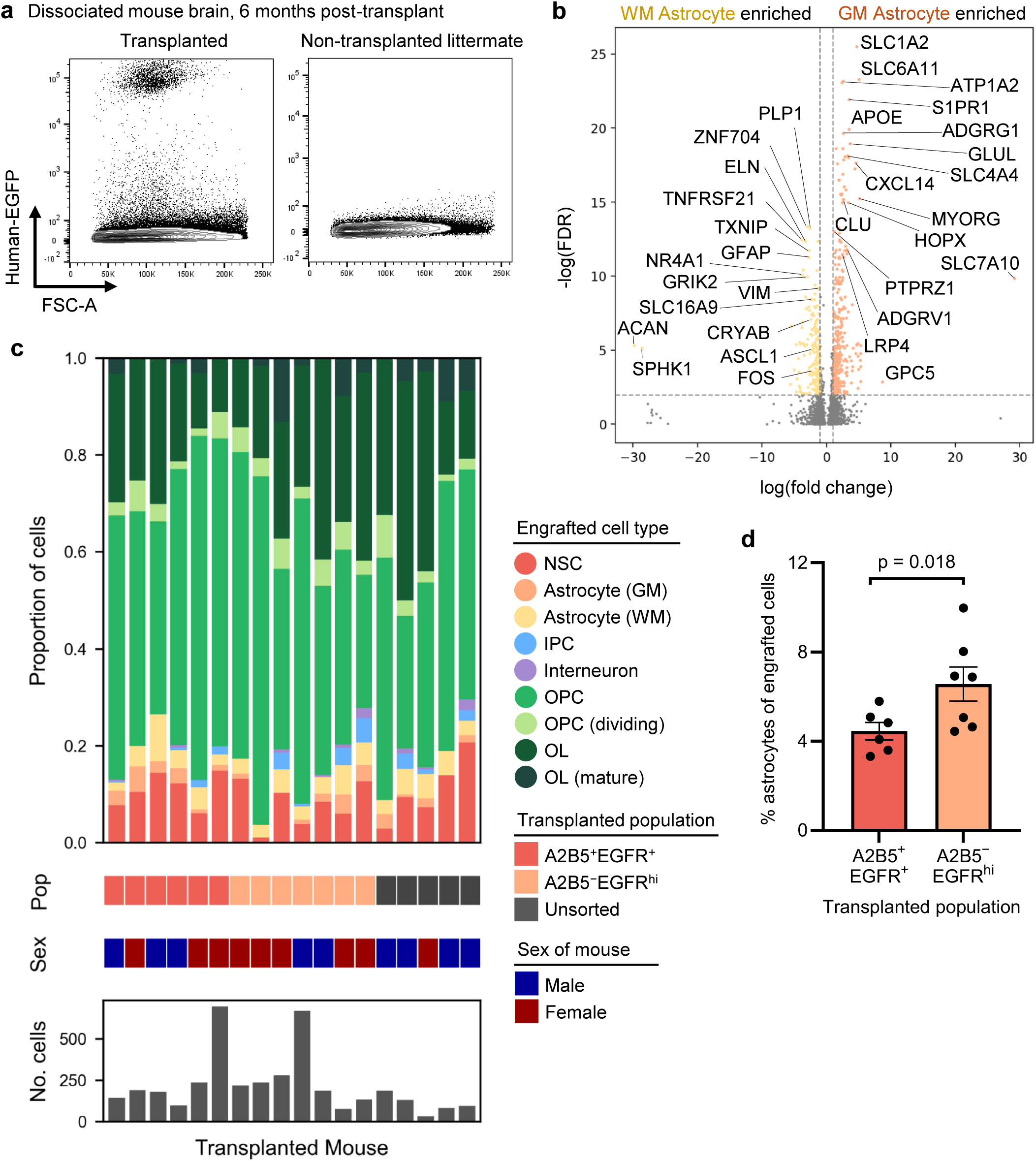
Analysis of engrafted postnatal human NSCs. **a,** Representative flow cytometry plots of dissociated mouse brains, 6 months post-transplant. Left, transplanted mouse; right, non-transplanted littermate. **b,** Volcano plot showing differentially-expressed genes between white matter astrocytes and grey matter astrocytes recovered from engrafted human cells. **c,** Stacked bar plot showing relative proportions of cell types recovered from each of 18 transplanted mice. Each bar represents one transplanted mouse. Below each bar shows the transplanted population, sex of the mouse, and number of human cells recovered from that mouse. **d,** Bar plot showing percent of astrocytes among engrafted cells, for mice transplanted with either A2B5^+^EGFR^+^ cells (left) or A2B5^−^EGFR^hi^ cells (right). Each dot represents one mouse.

**Extended Data Fig. 9:**
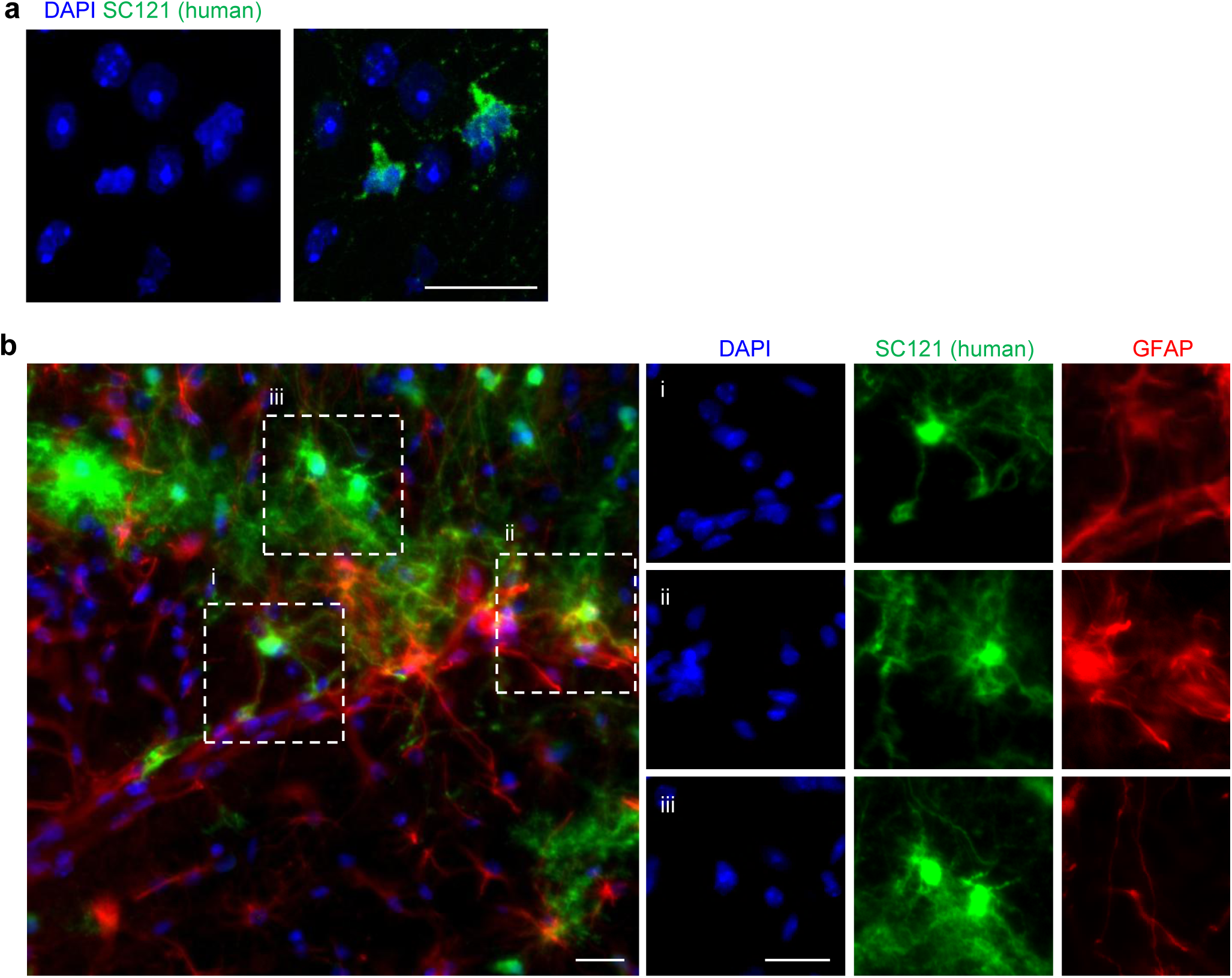
Histology of engrafted postnatal human NSCs. **a,** Immunofluorescence images demonstrating nuclear morphology of human cells (white arrowheads) engrafted in the mouse brain. **b,** Mouse subcortical white matter, stained for DAPI (blue), SC121 (green), and GFAP (red). Scale bars 20 μm.

**Extended Data Fig. 10:**
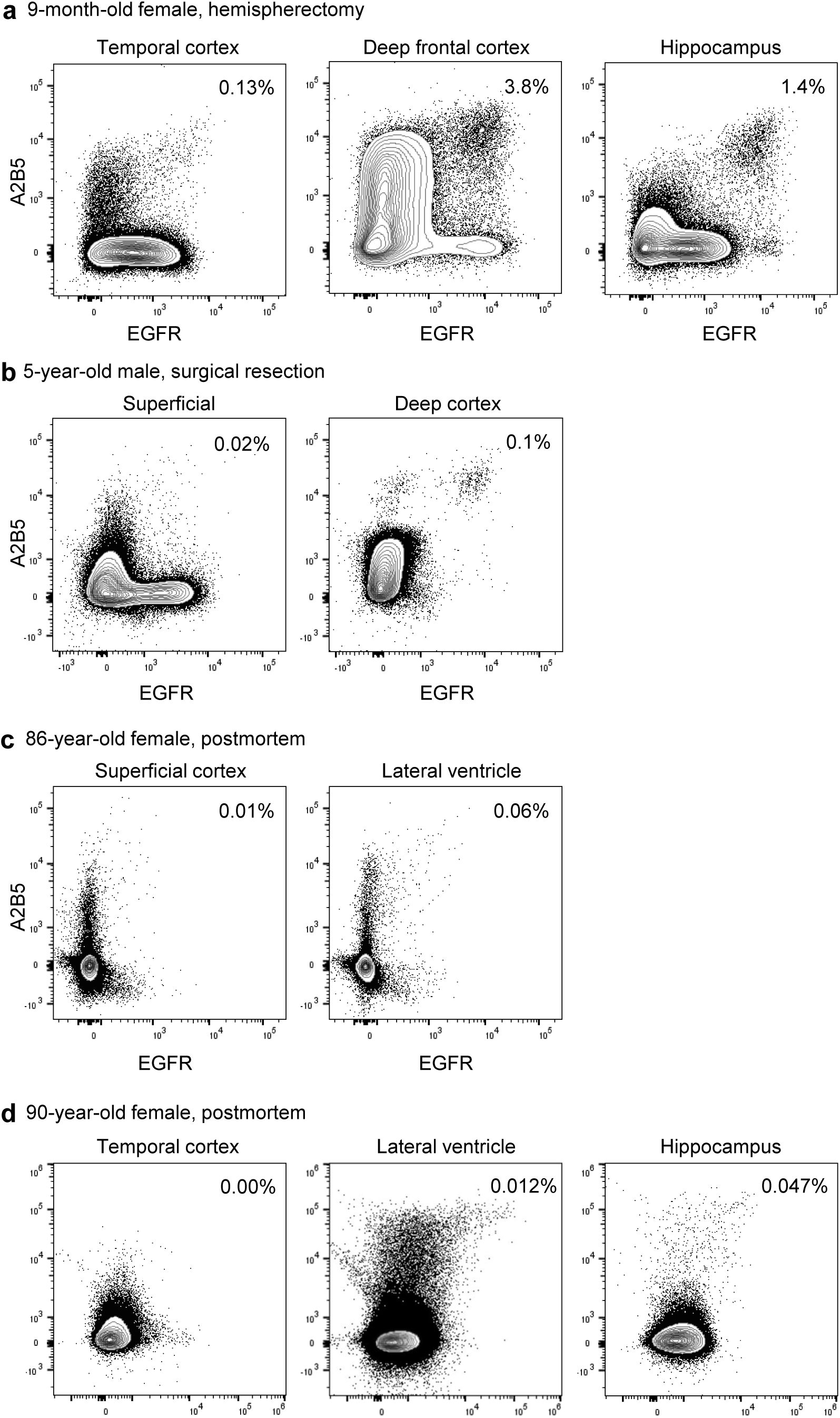
Immunophenotyping of human brain tissue across lifespan. Flow cytometry plots of dissociated postnatal human brain tissue of various anatomical origin. Events are pre-gated on live singlets of neural lineage. Frequency of A2B5^+^EGFR^+^ events denoted in top right corner.

